# Female resistance to the metabolic benefits of protein restriction is reversed by ovariectomy in mice

**DOI:** 10.64898/2026.03.31.715667

**Authors:** Bailey A. Knopf, Isaac Grunow, Brady Anderson, Tareq Rihawi, Michelle M. Sonsalla, Mariah F. Calubag, Reji Babygirija, Yang Liu, Fan Xiao, Chung-Yang Yeh, Dudley W. Lamming

## Abstract

Dietary protein intake mediates healthy aging in diverse species, with consumption of a low protein (LP) diet improving metabolic health in both humans and mice. In mice, the benefits of LP diets are sex-specific, with males exhibiting a stronger response to a LP diet than females. The reason for this sexually dimorphic response is unknown, but we hypothesized that sex hormones might be responsible for this difference. Here, we tested the role of sex hormones in the response to a LP diet by feeding intact and gonadectomized mice of both sexes either a Control (21% of calorie from protein) or LP (7% of calories from protein) diet, and assessing the effects on weight, body composition, glycemic control, and energy balance over the course of three months, followed by molecular and histological analysis of tissues from each group. We confirm that males show a stronger metabolic response to an LP diet than females, but that ovariectomy sensitizes female mice to the metabolic effects of an LP diet, making them respond more similarly to males; conversely, castration does not substantially impact the response of males to an LP diet. Molecularly, we find that gonadectomy and sex are important interactors that mediate the response of mechanistic target of rapamycin (mTOR) signaling, lipid homeostasis, and thermogenesis to an LP diet. Together, this data shows that the resistance of female mice to an LP diet is mediated by ovarian hormones and suggests the possibility that older female humans might receive enhanced benefits from LP diet feeding.

## Introduction

Over the past two decades, dietary protein has emerged as a critical regulator of metabolic health in species ranging from flies to mice and humans (Mair, Piper et al. 2005, Levine, Suarez et al. 2014, Solon-Biet, McMahon et al. 2014, Solon-Biet, Mitchell et al. 2015, Green, Lamming et al. 2022). Surprisingly, while high dietary protein intake is generally thought of as beneficial, a growing set of studies suggest that in humans, higher protein intake is associated an increased risk of age-related diseases, including diabetes (Sluijs, Beulens et al. 2010, Levine, Suarez et al. 2014, Ni Lochlainn, Bowyer et al. 2023). In randomized clinical trials, protein restriction, defined as reduction of dietary protein such that protein calories account for approximately 7-10% of total calories, improves human metabolic health, reducing weight and adiposity by boosting energy expenditure, and improving glycemic control (Fontana, Cummings et al. 2016, Ferraz-Bannitz, Beraldo et al. 2022, Nicolaisen, Lyster et al. 2025).

Laboratory studies support the idea that low protein (LP) diets are metabolically healthy, with multiple studies showing that LP diets have similar positive metabolic impacts on the health of rodents, reducing weight and adiposity, boosting energy expenditure, and improving glycemic control (Laeger, Henagan et al. 2014, Fontana, Cummings et al. 2016, Maida, Zota et al. 2016). However, all of the human randomized clinical trials discussed above and most of the rodent studies were conducted exclusively in males. While medical professionals advise individuals to consume 0.8-1.2 g/kg BW/day of protein regardless of sex (Nations 2007), studies in mice have found that the benefits of an LP diet are highly dependent on sex (Green, Pak et al. 2022).

Female mice exhibit a substantially reduced metabolic response to the effects of an LP diet compared to males (Larson, Russo et al. 2017, Green, Pak et al. 2022), however the mechanisms responsible for this blunted response remain undefined. One potential mechanism is FGF21, a hormone that regulates energy expenditure and stimulates food intake that has been shown to be required of many of the benefits of LP diets in males. One possibility for the blunted response is that an LP diet is less efficient at inducing FGF21 in females (Laeger, Henagan et al. 2014, Larson, Russo et al. 2017, Hill, Albarado et al. 2022).

Many sexually dimorphic responses are mediated in part by gonadal sex hormones (Garratt, Bower et al. 2017, Garratt, Lagerborg et al. 2018, Arriola Apelo, Lin et al. 2020). We hypothesized that gonadectomy would either blunt the metabolic responses of male mice to a LP diet or sensitize female mice to an LP diet. We therefore examined how mice responded to being fed either a Control (CTL, 21% of calories from protein) or low protein (LP, 7% of calories from protein) diet following gonadectomy. We found that while gonadectomy did not impact the response of male mice to an LP diet, ovariectomy sensitized female mice to the metabolic benefits of a LP diet.

Molecularly we show that sex, diet, and gonadectomy all independently alter lipolysis, lipogenesis, and thermogenesis. Similarly, these conditions alter mTOR signaling in the liver and skeletal muscle. This study highlights the important role that sex organs play in dietary intervention studies, and the necessity to continue exploring their role.

## Methods

### Animal care, Surgery, and Diet

All procedures were performed in accordance with institutional guidelines and were approved by the Institutional Animal Care and Use Committee of the William S. Middleton Memorial Veterans Hospital.

Male and female C57BL/6J mice, 7 weeks of age (n=12 per group), were obtained from The Jackson Laboratory (#000664, Bar Harbor, ME, USA). Prior to arrival at our facility, all male mice underwent either surgical sham or castration, and all female mice underwent either surgical sham or ovariectomy. These procedures were conducted under isoflurane anesthesia at 5-weeks of age by surgeons at The Jackson Laboratory. Briefly, the testis and the ovaries were excised, and the surgical incision was closed. Sham mice underwent the same surgical incision, the gonads were observed, and the incision was closed. Postoperatively, mice were examined daily by surgery technicians. Mice were monitored for 2 weeks prior to shipment.

Upon arrival, all mice were placed on a Teklad Global 18% protein rodent diet (Envigo 2018) for one week while they were acclimating to the animal research facility. All animals were housed in static microisolator cages in a specific pathogen free mouse facility with a 12:12 h light-dark cycle. Mice were dually housed and maintained at approximately 22°C with *ad libitum* access to water. Daily health checks were performed by animal staff.

At 8 weeks of age, mice were randomly assigned to a control 21% protein diet (TD. 181061) or a LP 7% protein diet (TD.10192) manufactured by Inotivo (formerly Envigo). Complete diet information is shown in **Table 1**. All animals were dually housed and monitored weekly for body weight and food consumption. Food consumption was calculated by measuring the difference between food put into the cage on Friday and the remaining food on Monday. Food consumption was normalized to body weight for each mouse. Every 3 weeks, mouse body composition was determined using an EchoMRI Body Composition Analyzer.

**Table 1.**
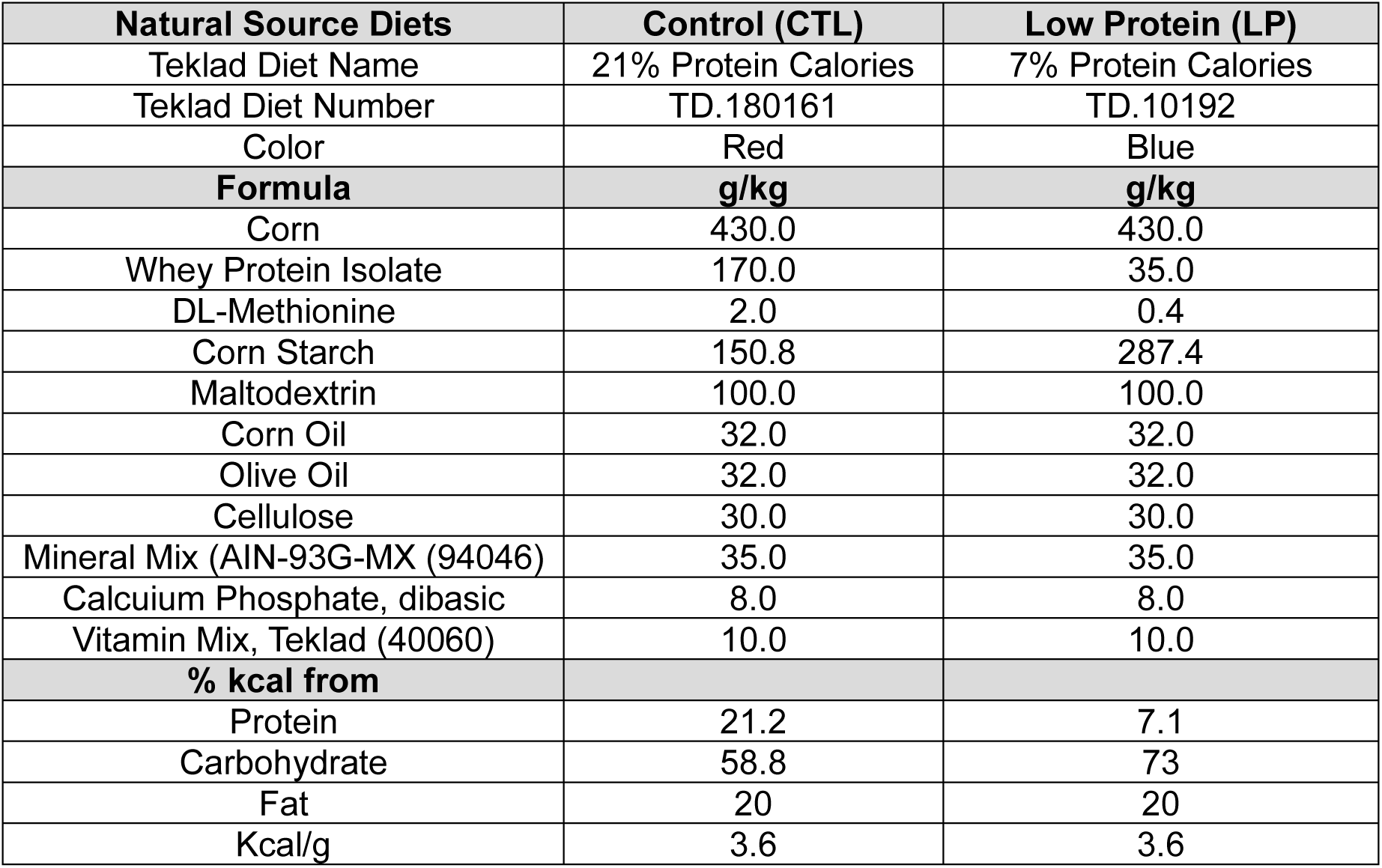
Dietary formulas used throughout the experiment.

### In vivo procedures

Glucose, insulin and alanine tolerance tests were performed after 3 weeks on the CTL or LP diet. For glucose and alanine tolerance tests the animals were fasted for 16 hours (overnight) while for insulin tolerance tests the animals were fasted for 4 hours. Mice were injected with either glucose (1 g/kg), insulin (0.75 U/kg), or alanine (2 g/kg) intraperitoneally (i.p.), and tail glucose measurements were taken using a Bayer Contour blood glucose meter (Bayer, Leverkusen, Germany) every 15 minutes for 90-120 minutes following injection. 16-hour fasted and 4-hour refed blood and glucose measurements for HOMA2IR was collected by tail vein.

After 6 weeks of dietary intervention, animals were temporarily housed in an Oxymax/CLAM-HC metabolic chamber system (Columbus Instruments) for approximately 48 hours. The first approximately 24 hours of data were discarded and served as an acclimation period for the animals; data collected during the subsequent 24 hours was then used for data analysis.

All animals were euthanized after a 16-hour overnight fast followed by 4 hours of refeeding the following morning. Terminal submandibular blood was collected into a tube containing EDTA, and animals were euthanized using cervical dislocation. Tissues were rapidly collected and flash frozen in liquid nitrogen. Blood was centrifuged at 2,000g for 5 minutes to isolate plasma. All plasma and tissues were stored at -80°C for future molecular analysis.

### Assays and Kits

Blood for fasting insulin was collected following a 16-hour overnight fast. Plasma insulin levels were determined using an ultra-sensitive mouse insulin ELISA kit (90080) from Crystal Chem (Elk Grove Village, IL, USA). Blood for circulating FGF21, testosterone, and estradiol levels was collected from submandibular bleeding immediately prior to euthanasia; plasma was assayed with a mouse/rat FGF21 quantikine ELISA kit (MF2100) from R&D Systems (Minneapolis, MN, USA), and testosterone (582701) and estradiol (501890) quantikine ELISA kits from Caymen Chemicals (Ann Arbor, MI, USA). Liver and plasma triglycerides and cholesterol were measured using Triglyceride Colorimetric Assay Kit (Pointe Scientific, #C75101L) and Cholesterol Colorimetric Assay Kit (Pointe Scientific, T7532500) following the manufacturer’s protocol.

### Quantitative PCR

Liver RNA was extracted using TRI Reagent (Sigma, St Louis, MO, USA) following the manufacturer’s protocol. RNA concentration and purity were determined using a Nanodrop (Thermo Fisher Scientific) measuring the absorbance at 260/280 nm. 1 μg of RNA was used to generate cDNA (Superscript III; Invitrogen, Carlsbad, CA, USA). Oligo dT primers and primers for real-time PCR were obtained from Integrated DNA Technologies (IDT, Coralville, IA, USA); *Atgl* (F 5’-ATATCCCACTTTAGCTCCAAGG-3’, R 5’-CAAGTTGTCTGAAATGCCGC-3’), *Lipe* (F 5’-CTGAGATTGAGGTGCTGTCG-3’, R 5’-CAAGGGAGGTGAGATGGTAAC-3’), *Fasn* (F 5’-CCCCTCTGTTAATTGGCTCC-3’, R 5’-TTGTGGAAGTGCAGGTTAGG-3’), *Scd-1* (F 5’-AGAAGGTGCTAACGAACAGG-3’, R 5’-CTGACCTGAAAGCCGAGAAG-3’), *Fgf21* (F 5’-CAAATCCTGGGTGTCAAAGC-3’, R 5’-CATGGGCTTCAGACTGGTAC-3’), *Gpt1* (F 5’-CCACTCAGTCTCTAAGGGCTAC-3’, R 5’- ACACAACCGCACGCTCATCAGT-3’), *Dgat1* (F 5’-TGGTGTGTGGTGATGCTGATC-3’, R 5’ -GCCAGGCGCTTCTCAA-3’), *Acc1* (F 5’ - AAGGCTATGTGAAGGATG-3’, R 5’ -CTGTCTGAAGAGGTTAGG-3’), *Ucp1* (F 5’ - GCATTCAGAGGCAAATCAGC-3’, R 5’ -GCCACACCTCCAGTCATTAAG-3’) *Cidea* (F 5’ - GAATAGCCAGAGTCACCTTCG-3’, R 5’ -AGCAGATTCCTTAACACGGC-3’), *Elovl3* (F 5’ - ATGCAACCCTATGACTTCGAG-3’, R 5’ -ACGATGAGCAACAGATAGACG-3’), *Ckb* (F 5’ - GCCTCATCTAGATCGAAACTC-3’, R 5’ -GGCATGTGAGGATGTAGCCC-3’) and *Bact (*F 5’-GATGTATGAAGGCTTTGGTC-3’, R 5’- TGTGCACTTTTATTGGTCTC-3’). Reactions were run on a StepOne Plus machine (Applied Biosystems, Foster City, CA, USA) with Sybr Green PCR Master Mix (Invitrogen). Actin was used to normalize the results from gene-specific reactions.

### Western Blotting

Liver and skeletal muscle tissue was homogenized in cold RIPA buffer supplemented with EDTA-free Protease and Phosphatase Inhibitor Mini Tablet (Thermo Fisher Scientific, A32961) as previously described (Richardson, Konon et al. 2021, Yeh, Chini et al. 2024, Calubag, Ademi et al. 2025), using a FastPrep 24 (M.P. Biomedicals, Santa Ana, CA, USA) with screw top microcentrifuge tubes (2682558, Fisher Scientific, Waltham, MA, USA) and zirconium ceramic oxide beads (10158-552, VWR, Radnor, PA). Protein lysates were centrifuged at 13,300 rpm for 10 minutes and supernatant was collected. Protein concentration was determined by Bradford Assay (Pierce Biotechnology, Waltham, MA, USA). 20 μg of protein lysate was separated by SDS-PAGE (sodium dodecyl sulfate-polyacrylamide gel electrophoresis) using a 16% Novex Tris-Glycine Gel (Thermo Fisher Scientific). Protein was transferred onto a PVDF membrane (EMD Millipore, Burlington, MA, USA) and blocked for 20 minutes in 5% milk in 1X TBST at room temperature. Membranes were incubated overnight in primary antibody (in 5% milk and all phosphoresidues in 5% BSA) at 4°C with gentle rocking and subsequently incubated with HRP goat anti-rabbit secondary antibody (Cell Signaling Technology #7074, 1:2,000 in 5% milk) for 1 hour at room temperature. Membranes were imaged using a Bio-Rad Chemidoc MP imaging station (Bio-Rad, Hercules, CA, USA). Primary antibodies were used at 1:1,000 and were purchased from Cell Signaling Technology (Danvers, MA, USA): p-T389 p70 S6K1 (#9205), p70 S6K1 (#2708), p-Ser240/244 S6 (#2215), S6 (#2217), p-Thr37/46 4E-BP1(#2855), 4E-BP1 (#9452), p-S51 eiF2α (#3398), eiF2α (#5324), p- S473 AKT (#4060), p-T308 AKT (#9275), AKT (#4691), HSP90 (#4877). Imaging was performed using a Bio-Rad Chemidoc MP imaging station (Bio-Rad, Hercules, CA, USA). Quantification was performed by densitometry using ImageJ software.

### Liver histology

After euthanasia, a portion of each liver was placed into optimal cutting temperature compound (OCT) and froze on dry ice. Samples were sectioned and stained with Oil Red O by the UWCCC Experimental Animal Pathology Laboratory. Images of the stained liver were taken using an EVOS microscope (Thermo Fisher Scientific Inc., Waltham, MA, USA) at a magnification of 40X. For quantification of lipid droplet size, 9 independent images were taken from each sample and quantified using ImageJ by a blinded investigator.

### Statistical Analysis

Most statistical analysis were performed using Prism, version 10 (GraphPad Software, San Diego, CA, USA). Data are presented as the mean ± SEM unless otherwise specified. Tests involving multiple factors were analyzed by a two-way or three-way analysis of variance (ANOVA) with sex, diet, and gonadectomy as categorial variables followed by Šidák’s multiple comparison *post hoc* testing for pair-wise comparisons. Energy expenditure differences were detected using analysis of covariance (ANCOVA). ANCOVA analysis assumes a linear relationship between the variables and their covariates. If the slope is equal between the groups, then the regression lines are parallel, and elevation is tested to determine any differences (i.e., if the slopes are statistically significantly different, elevation will not be determined). PCA plots were produced utilizing R, version 4.3.0, by imputing missing data and scaling the data using the package “missMDA” (Josse 2016). Visualization of the PCA plots were performed using the following packages: “FactoMineR” (Lê 2008), “Factoextra” (Kassambara 2020), and “corrplot” (Wei 2024). Data distribution was assumed to be normal, although not formally tested. Statistical parameters can be found in each figure and figure legend. In all figures, n represents the number of biologically independent animals.

## Results

### The response to a 12-week LP dietary intervention on weight and body composition is dependent on biological sex and the presence of sex organs

We gonadectomized wild-type C57BL/6J mice at 5 weeks of age; sham surgeries were performed on mice of both sexes to control for any effects of undergoing surgery. Mice of each sex and surgical status were then randomized at 8 weeks of age to two diet groups: Control (CTL, 21% calories from protein) or a low protein (LP, 7% calories from protein) (**Fig. 1A**). These two diets are isocaloric, with 20% of calories from fat, and carbohydrates were used to replace the calories from protein in the LP diet (**Fig. 1B**). Mice were maintained on these diets for 12 weeks prior to euthanasia. After euthanasia, we analyzed plasma testosterone and estradiol levels to confirm that gonadectomy robustly reduced the sex hormone levels (**Figs. S1A-B**).

**Figure 1.**
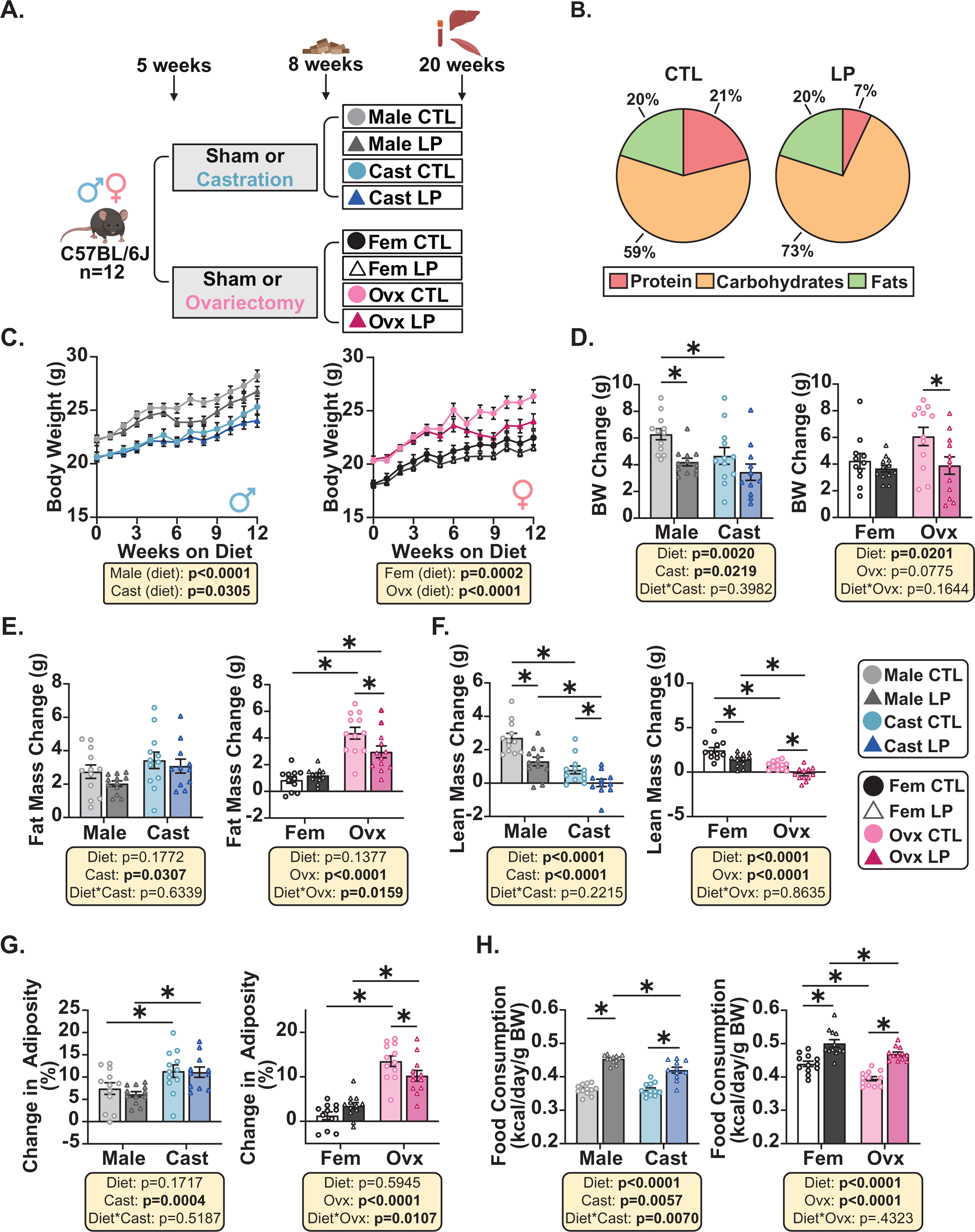
Changes in body composition are dependent on sex, gonadectomy, and dietary condition. (A) Experimental schematic: male and female C57BL/6J mice underwent either gonadectomy or surgical sham at 5 weeks of age. Mice were then randomly assigned to a Control (CTL) protein diet or a Low Protein (LP) diet at 8 weeks of age. Mice were euthanized after 12 weeks of dietary intervention and tissues were collected. (B) Macronutrient composition of the diets. (C). Body weights throughout the dietary intervention period. (D) Quantification of the change in body weight during the 12-week study. (E-G) Change in fat mass (E), lean mass (F), and adiposity (G) by the end of the 12-week experiment. (H) Average food intake measured weekly and calculated per day normalized to body weight over the 12 weeks. (C-H) n=11-12 mice per group. (C-H) Statistics for the overall effects of diet, gonadectomy, and the interaction represent the p value from a two-way ANOVA. *p<0.05 from a Šidák’s post-test examining the effect of parameters identified as significant in the two-way ANOVA. Data are represented as mean ±SEM. Abbreviations: CTL (21% control protein diet), LP (7% low protein diet), Cast (castration), Fem (female), Ovx (ovariectomy). Created in BioRender. Knopf, B. (2026) https://BioRender.com/bibd348

Prior to starting the experiment, we observed that gonadectomy significantly affected the body weight of male and female mice, with castrated (Cast) mice weighing significantly less and ovariectomized (Ovx) mice weighing significantly more than their Sham surgery counterparts (Male and Female (Fem), respectively) (**Fig. S1C**). Surprisingly, the increase in body weight for Ovx mice prior to dietary intervention was driven by an increase in lean mass (**Figs. S1D-E**). During the 12-week dietary intervention, all mice gained weight irrespective of sex, diet, or gonadectomy (**Figs. 1C-D**). Similar to what we have previously reported (Green, Pak et al. 2022), there was an overall effect of diet on male mice, with both Male and Cast LP-fed animals weighing less than their CTL-fed counterparts (p<0.0001 and p=0.0305, respectively) (**Figs. 1C-D**). While there was also an overall significant effect of diet on female mice, this was driven by a significant effect of a LP diet on the body weight of Ovx mice, and there was no significant difference in body weight between CTL and LP-fed female mice at the conclusion of the experiment (**Fig. 1D**).

Consistent with previous work (Green, Pak et al. 2022), we observed a sex-specific response of fat mass gain on a LP diet. Overall, we observed a significant sex and gonadectomy effect of fat mass change over the 12-weeks (**Fig. 1E**). Castrated mice do not exhibit the approximately 20% reduction in fat mass that is canonically seen in intact C57BL6/J male mice on a LP diet (**Fig. 1E, S1F**). Interestingly, Ovx LP-fed mice showed decreased fat mass of around 25% that was not observed in the intact female counterparts (**Figs. 1E, S1F**). All LP-fed mice regardless of sex or gonadectomy exhibit significantly reduced lean mass compared to their CTL-fed counterparts (**Figs. 1F, S1G**). Ovx LP-fed mice were the only group to exhibit significantly reduced adiposity after the 12-week intervention when compared to the CTL-fed mice (**Figs. 1G, S1H**).

Despite the decreased weight gain of LP-fed mice, we found that all LP-fed mice, regardless of sex or surgical status, had increased average food consumption during the 12-week intervention (**Fig. 1H**). We observed a significant effect of surgery in both sexes as well as a significant diet x surgery interaction in males, which we interpret as a LP diet stimulating food intake less in Cast mice- Ovx mice have a lower overall food intake normalized to body weight regardless of diet, likely due in part to the greater weight of Ovx mice **(Fig. 1H**).

### Sex, gonadectomy, and diet all impact energy balance in mice

To gain insight into how a LP diet reduces body weight and adiposity in the absence of calorie restriction, we assessed energy balance using metabolic chambers to measure energy expenditure (EE) via indirect calorimetry. Consistent with previous data (Laeger, Henagan et al. 2014, Hill, Laeger et al. 2017, Green, Pak et al. 2022), we found that an LP diet increases EE during the dark cycle across all groups regardless of sex or gonadectomy, even when correcting for differences in body weight through ANCOVA (**Figs. 2A-B**). We also observed an overall effect of surgery on dark phase EE in both sexes, with decreased energy expenditure in both Cast and Ovx mice relative to intact controls. We observed similar results during the light cycle, with LP diets robustly increasing EE across sexes and surgical conditions (**Fig. S2A**).

**Figure 2.**
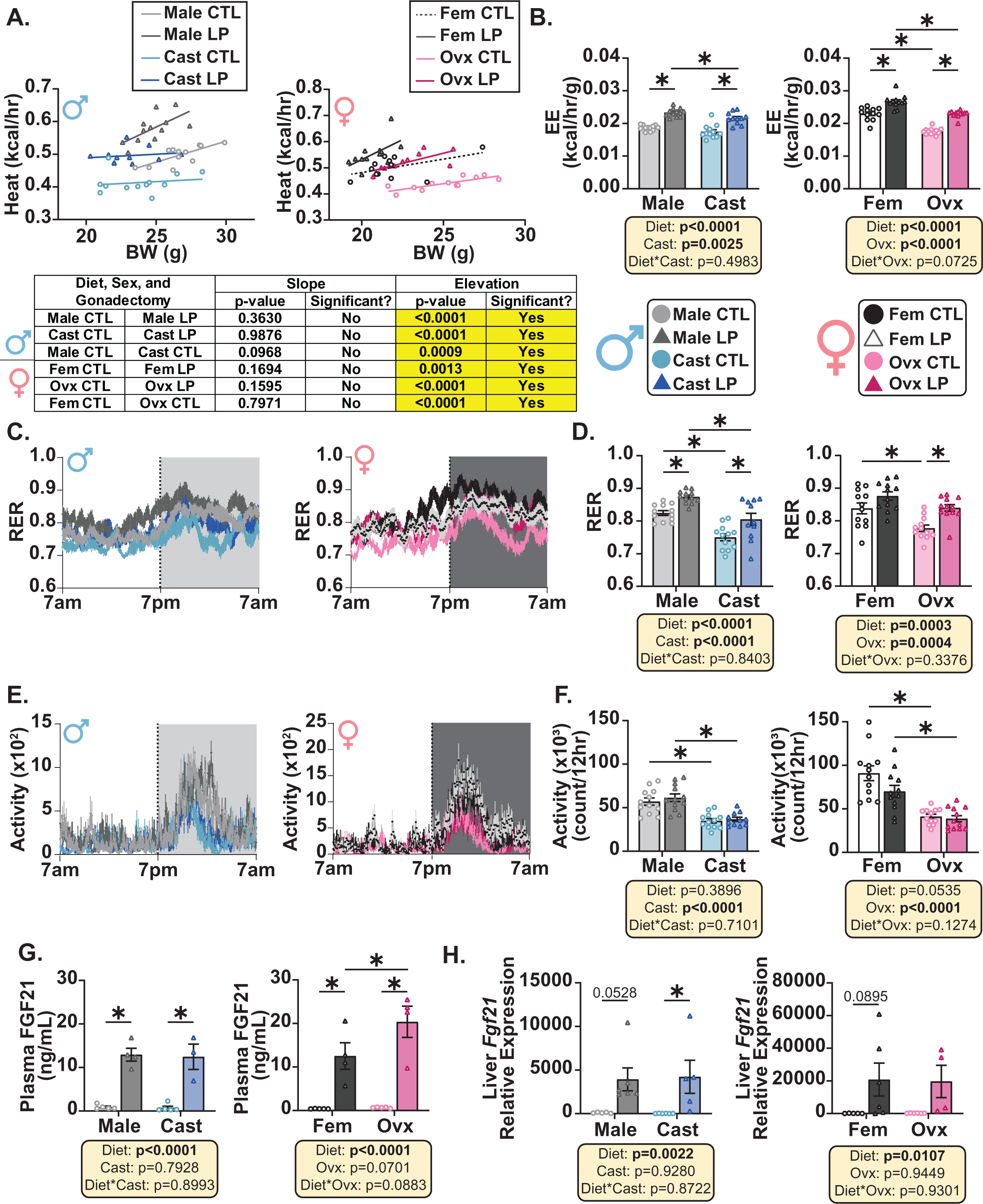
The impact of sex, diet, and gonadectomy on energy balance during the dark cycle. (A) ANCOVA of energy expenditure (EE) with body weight as a covariate during the 12-hour dark cycle. (B) Average EE during the dark cycle normalized to body weight. (C) Respiratory Exchange Ratio (RER) over a 24-hour period. (D) Average RER during the dark cycle. (E) Spontaneous activity over a 24-hour period. (F) Average spontaneous activity during the dark cycle, calculated as laser beam breaks. (G) Circulating FGF21 expression quantified from refed plasma using ELISA. (H) The mRNA expression of *Fgf21* in the liver of mice. (A-F) n=11-12 mice per group, (G-H) n=3-5 mice per group. (A) Data for each individual mouse is plotted, simple linear regression (ANCOVA) was calculated to determine if the slopes or elevations are equal. (B, D, F-H) Statistics for the overall effects of diet, gonadectomy, and the interaction represent the p value from a two-way ANOVA. *p<0.05 from a Šidák’s post-test examining the effect of parameters identified as significant in the two-way ANOVA. Data are represented as mean ±SEM. Abbreviations: CTL (21% control protein diet), LP (7% low protein diet), Cast (castration), Fem (female), Ovx (ovariectomy), BW (body weight), EE (energy expenditure), RER (respiratory exchange ratio), FGF21 (fibroblast growth factor 21).

We analyzed the Respiratory Exchange Ratio (RER) over a 24-hour period and found that LP diets significantly increase RER regardless of sex or gonadectomy (**Figs. 2C-D**). Increased RER suggests a shift in fuel utilization towards carbohydrates and away from fats, so it is expected that LP diets containing greater proportions of carbohydrates will result in increased RER. In addition, there was an overall effect of surgery on RER in both sexes, with Cast and Ovx mice having reduced RER relative to intact mice, suggesting an increased utilization of fats during the dark cycle (**Fig. 2D**). During the light cycle, RER is overall reduced across groups, however we still observe the same diet and gonadectomy effect that is observed in the dark cycle (**Fig. S2B**). While we observed no effect of a LP diet on spontaneous activity during the dark cycle, we did see a strong effect of gonadectomy, with reduced spontaneous activity in both Cast and Ovx mice relative to intact controls (**Fig. 2E-F**). These results are not as apparent in the light cycle (**Fig. S2C**), as there is only an Ovx effect of reduced activity compared to intact females. Physical activity increases RER suggesting that the decrease in RER seen with gonadectomized mice may be due to decreased activity (Simonson and DeFronzo 1990). This decrease in activity suggests that the reduction in body weight observed with an LP diet is not due to activity, but rather due to remodeling of metabolic pathways.

The hormone FGF21 is well established to mediate many of the metabolic phenotypes observed with dietary PR (Laeger, Henagan et al. 2014, Hill, Albarado et al. 2022). Regardless of sex or gonadectomy, we observed LP diets significantly increase FGF21 levels in the plasma (**Fig. 2G**). The increase in plasma levels of FGF21 is likely driven by the significant upregulation of *Fgf21* transcription in the liver of LP fed mice (**Fig. 2H**).

### Ovariectomy sensitizes mice to LP-induced improvements in glycemic control

To examine the interaction between gonadectomy and diet on glucose homeostasis, we performed a series of assessments, including glucose, insulin, and alanine tolerance tests and collecting blood to measure fasting glucose and insulin. As expected, we found that an LP diet significantly improved glucose tolerance in intact males but did not significantly improve glucose tolerance in intact females (**Figs. 3A-B**). While castration blunted the impact of an LP diet on glucose tolerance, there was not a significant diet x surgery interaction in males; in contrast, there was a diet x surgery interaction in females, with Ovx mice showing a significant improvement in glucose tolerance on a LP diet (**Figs. 3A-B**).We found that gonadectomy itself also impacts glucose tolerance in a sexually dimorphic manner. Cast mice exhibit improved glucose tolerance compared to their intact male counterparts, and Ovx mice exhibit impaired glucose tolerance compared to intact females (**Figs. 3A-B**). An LP diet improves insulin tolerance in both males and females, with no diet x surgery interaction in either sex (**Figs. 3C-D**). Improved glucose tolerance can result from improved suppression of hepatic gluconeogenesis or enhanced insulin secretion. We therefore performed an alanine tolerance test; alanine is a hepatic substrate for gluconeogenesis (Mutel, Gautier-Stein et al. 2011). In all groups, LP feeding improved alanine tolerance; there was also a significant effect of surgery in female mice, with Ovx mice exhibiting worse alanine tolerance than Sham controls (**Figs. S3A-B**).

**Figure 3.**
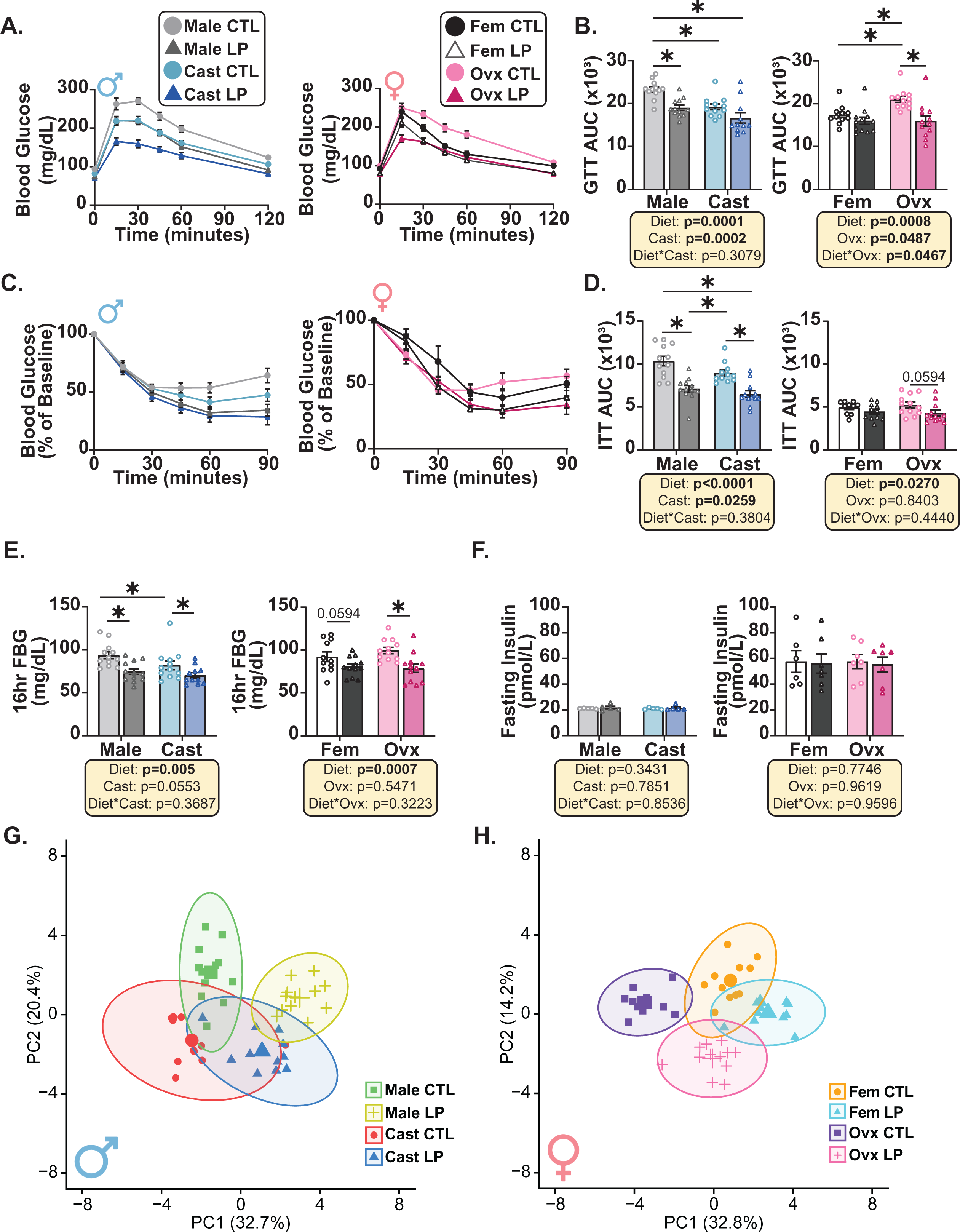
Ovx LP mice mirror male LP mice on measures of glucose homeostasis. (A-B) Glucose tolerance test (GTT) and area under the curve (AUC) after 8 weeks on diet. (C-D) Insulin tolerance test (ITT) normalized to baseline glucose values and calculated AUC. (E) Fasting blood glucose (FBG) during a 16-hour fast. (F) Fasting plasma insulin levels were determined using ELISA. (G-H) PCA on phenotypic measurements from male (G) or female (H) mice indicating separation between the groups along PC1 and PC2. (A-E, G-H) n=11-12 mice/group. (F) n=4-7 mice/group. (B, D, E-F) Statistics for the overall effects of diet, gonadectomy, and the interaction represent the p value from a two-way ANOVA. *p<0.05 from a Šidák’s post-test examining the effect of parameters identified as significant in the two-way ANOVA. Data are represented as mean ±SEM. Abbreviations: CTL (21% control protein diet), LP (7% low protein diet), Cast (castration), Fem (female), Ovx (ovariectomy), GTT (glucose tolerance test), AUC (area under the curve), ITT (insulin tolerance test), FBG (fasting blood glucose).

To further analyze the effect of sex organs and the response to a LP diet on glucose homeostasis, we collected fasting blood and determined fasting blood glucose and insulin levels (**Figs. 3E-F**). We observed a significant effect of diet on fasting blood glucose levels in both sexes, with LP reducing fasting blood glucose in all groups. There was no effect of diet or surgery on fasting insulin (**Figs. 3E-F**). In addition, all female mice, regardless of surgery status, had significantly higher fasting insulin levels than males (**Fig. 3F**).

Finally, we used the fasted blood glucose and insulin levels to calculate insulin resistance using homeostasis model assessment (HOMA2-IR) (Levy, Matthews et al. 1998). All female mice had a higher HOMA2-IR than males due to their higher fasting insulin levels; we also observed an overall effect of diet on HOMA2-IR in female mice (**Fig. S3D**). We observed an overall effect of diet on beta cell function (HOMA %B), with a significant diet x surgery interaction in males; the LP diet boosted HOMA %B in both intact males and in Ovx females, but not in Cast males or intact Females (**Figs. S3E**).

To understand the contributions of various phenotypic outcomes between surgical conditions and diet, we performed a principal component analysis (PCA) using only the phenotypic data. We found that both sex and gonadectomy influenced the phenotypic outcome of a LP diet (**Figs. 3G-H**). Notably, in intact males and Ovx females, CTL and LP-fed mice do not overlap; conversely, CTL and LP-fed mice do overlap in Cast males and intact females. Thus, the overall response to an LP diet is greatest in intact males and Ovx females (**Figs. 3G-H**). For both sexes, the greatest contributors to PC1 are energy expenditure, food consumption, and RER (**Figs. S3F-G**). The greatest contributors for PC2 in the males are GTT AUC, activity, and lean mass change, while the females’ greatest contributors are changes in lean mass, 4-hour FBG, and activity (**Figs. S3F-G**).

### LP diets increase hepatic lipid storage regardless of sex and gonadectomy

Previous studies in rodents show that consuming an LP diet leads to an accumulation of lipids in the liver and ultimately can promote hepatic steatosis (Arvidsson Kvissberg, Hu et al. 2022). To assess how sex and gonadectomy alter lipid droplet composition in the liver, we performed Oil Red O (ORO) staining (**Fig. 4A**). Quantitative analysis revealed a significant diet effect of increased lipid droplet size and total staining area in LP-fed mice (**Figs. 4B-C**). We also observed a sex effect of total staining area with females having increased lipid staining compared to males (**Fig. 4C**).

**Figure 4.**
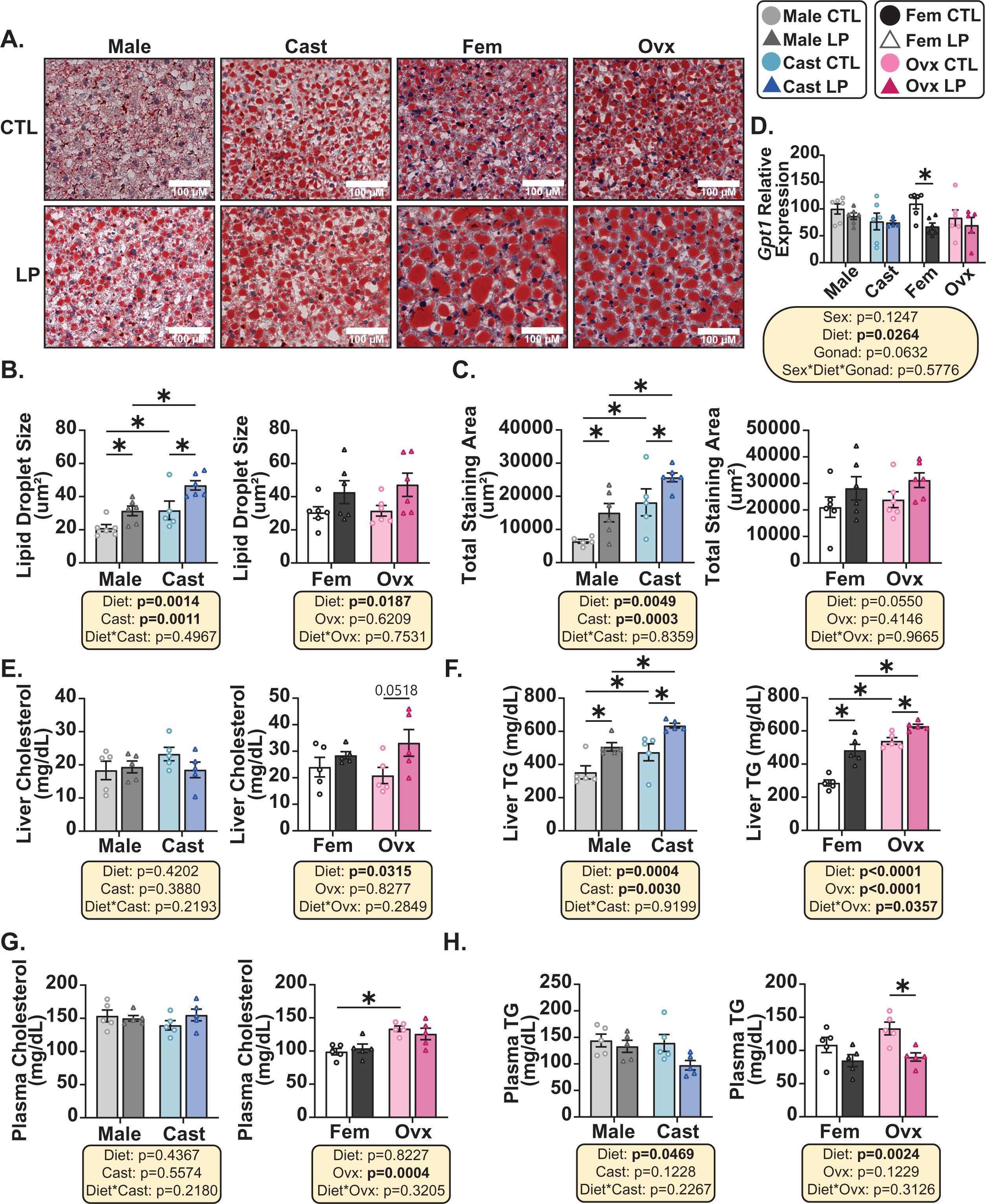
Characterization of hepatic and circulating lipid homeostasis with sex, diet, and gonadectomy. (A-C) Oil Red O (ORO) images of livers that were embedded in optimal temperature compound (OCT) and stained (representative images; scale bar= 100 μM, 40X magnification). Lipid droplet size (B) and overall staining area (C) were quantified for each image using ImageJ. (D) Quantification of *Gpt1* expression in the liver using qPCR. (E-F) Hepatic cholesterol (E) and triglycerides (TG) (F) quantification using colorimetric assays. (G-H) Circulating cholesterol (G) and TG (H) quantification on refed plasma. (A-H) n=5-6 mice/group. (B-C, E-H) Statistics for the overall effects of diet, gonadectomy, and the interaction represent the p value from a two-way ANOVA. *p<0.05 from a Šidák’s post-test examining the effect of parameters identified as significant in the two-way ANOVA. (D) Three-way ANOVA between sex, diet, and gonadectomy with post hoc Šidák’s adjusted test for pairwise comparisons, *p<0.05. p values for the overall effect of sex, diet, and gonadectomy and the interactions represent the significant p values from the three-way ANOVA. Data are represented as mean ±SEM. Abbreviations: CTL (21% control protein diet), LP (7% low protein diet), Cast (castration), Fem (female), Ovx (ovariectomy), ORO (oil red O), TG (triglycerides).

To determine if the increase in lipids present in the liver is detrimental to liver integrity, we analyzed hepatic glutamate-pyruvate transaminase (*Gpt1*) expression (Sookoian, Castano et al. 2016, Okun, Rusu et al. 2021). *Gpt1* transcription is associated with fatty liver disease in human patients (Sookoian, Castano et al. 2016), and acts as a marker for hepatic cellular health. While we observe changes in hepatic lipid homeostasis, we found a reduction in *Gpt1* expression upon dietary protein restriction (**Fig. 4D**), suggesting reduced dietary protein improves hepatic cellular health.

Because we observed increased accumulation of lipids in the liver, we quantified triglyceride (TG) and cholesterol content of the liver. For the males, we found no change in liver cholesterol levels regardless of gonadectomy (**Fig. 4E**). However, in the females we found a significant diet effect of LP increasing hepatic cholesterol (**Fig. 4E**). Consistent with the ORO staining, we observed a significant increase in liver TG in LP-fed mice regardless of sex or surgical condition (**Fig. 4F**), suggesting that the increased lipid droplets observed in ORO may be driven by increased triglycerides. We further found that the presence of sex organs reduces hepatic TG, suggesting an interaction between sex organs and hepatic lipid homeostasis (**Fig. 4F**). We next quantified plasma cholesterol levels and found that an LP diet did not alter plasma cholesterol levels, which is consistent with what we observed in the liver (**Fig. 4G**). With an increase in liver TG, we hypothesized that there may be an increase in circulating triglycerides as well, however we observed the opposite. Mice consuming an LP diet had reduced plasma TG levels, with Ovx mice exhibiting a significant reduction (**Fig. 4H**).

### The nutrient-sensing regulator, mTOR, is altered with gonadectomy, sex, and dietary protein content

Previous work in our laboratory has found tissue- and sex-specific differences in the master nutrient regulator, mechanistic Target Of Rapamycin (mTOR). Previous work reports sexual-dimorphism in mTOR complex 1 and complex 2 (mTORC1 and mTORC2) signaling, with higher activity in the livers of female mice (Baar, Carbajal et al. 2016). Studies from several laboratories have found that dietary protein regulates hepatic mTORC1 activity, as well as the activity of mTORC1 in other tissues (Harputlugil, Hine et al. 2014, Solon-Biet, McMahon et al. 2014, Lamming, Cummings et al. 2015). A recent report found that dietary protein, not insulin signaling, regulates mTORC1 induction by feeding in the liver (Kalafut, Cormerais et al. 2025). To determine the effect of an LP diet on components of the mTROC1 pathway in a gonadectomy and sex-specific manner, we analyzed the liver and muscle tissue of these mice.

As shown in **Figs. 5A-G**, we observed no significant diet, sex, or gonadectomy effect on phosphorylated S6 (**Fig. 5B**), phosphorylated 4E-BP1 (**Fig. 5C**), and phosphorylated eIF2α (**Fig. 5E**) in the liver. We saw significantly blunted phosphorylation of S6K1 in the livers of LP-fed mice, consistent with reduced mTORC1 activity (**Fig. 5D**). We found a sex and gonadectomy effect on phosphorylated S6K, with females having higher levels, and gonadectomized mice having lower levels (**Fig. 5D**). We observed an overall significant decrease in phosphorylated (S473) AKT due to gonadectomy, suggesting a potential impact of the sex organs on hepatic mTORC2 activity (**Fig. 5F**).

**Figure 5.**
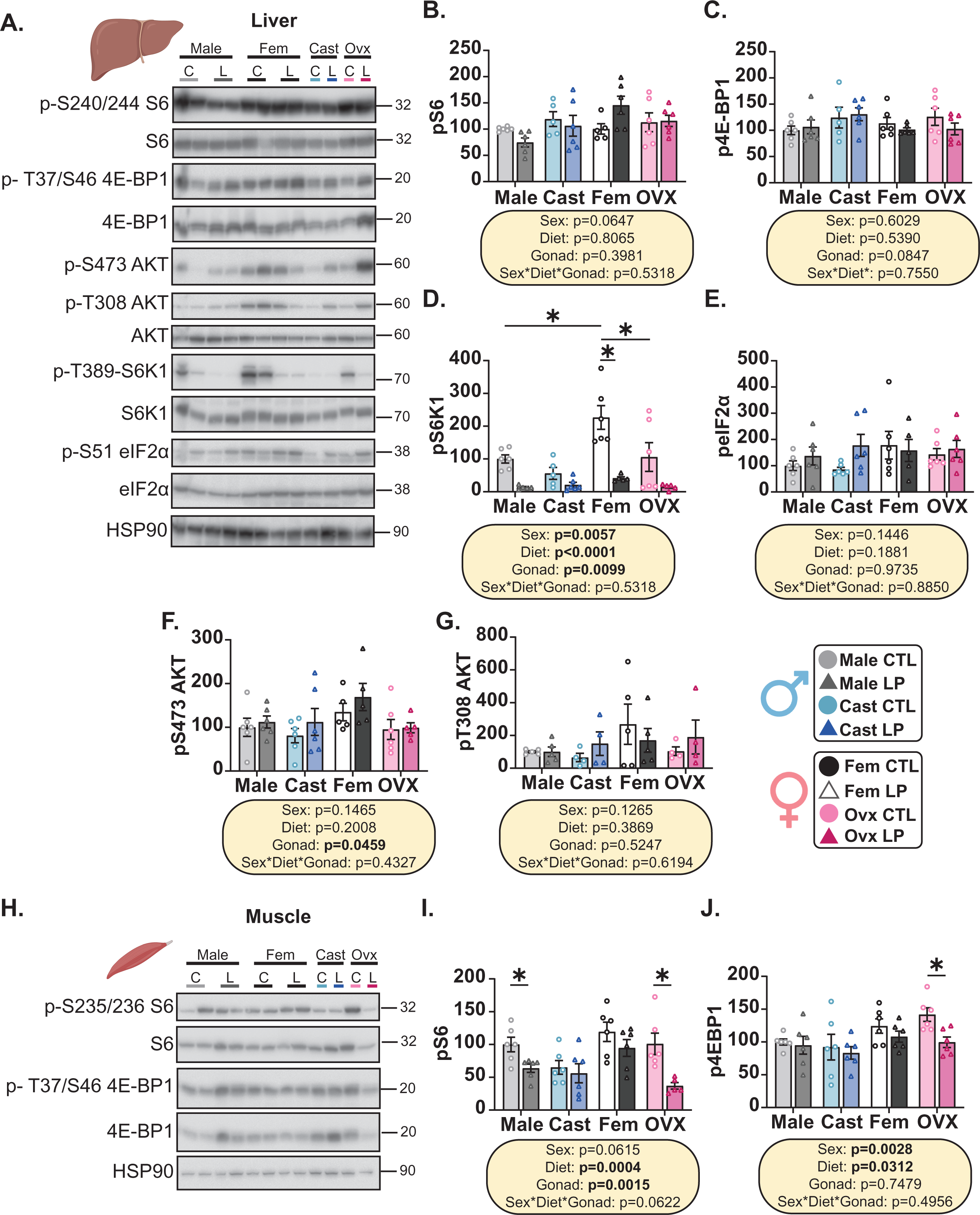
Tissue specific alterations in mTORC1 and mTORC2 activity with sex, gonadectomy, and diet. (A-G) mTOR signaling in the liver was determined by Western blot analysis of the following phosphorylated and unphosphorylated proteins; S6 (B), 4e-BP1 (C), S6K1 (D), eiF2α (E), AKT (F-G), HSP90 (loading control). (H-J) mTORC1 signaling in the muscle was determined by Western blot analysis of phosphorylated S6 (I), phosphorylated 4e-BP1 (K). HSP90 was used as a loading control. (B-G, I-J) Quantification was determined by normalizing phosphorylated protein to normal protein. (A-J) n=5-6 mice/group, quantification was determined using ImageJ by normalizing the phosphorylated protein to the unphosphorylated form. Three-way ANOVA between sex, diet, and gonadectomy with post hoc Šidák’s adjusted test for pairwise comparisons, *p<0.05. p values for the overall effect of sex, diet, and gonadectomy and the interactions represent the significant p values from the three-way ANOVA. Data are represented as mean ±SEM. Abbreviations: C, CTL (21% control protein), L, LP (7% low protein), Cast (castration), Fem (female), Ovx (ovariectomy), Gonad (gonadectomy). Created in BioRender. Knopf, B. (2026) https://BioRender.com/i0gripn

We wanted to determine how another metabolically active tissue, such as the skeletal muscle, exhibited changes in mTORC1 signaling dependent on diet, sex, and gonadectomy. We observed a significant effect of diet on the phosphorylation of both S6 and 4E-BP1 in skeletal muscle. (**Figs. 5H-J**). Female mice have significantly increased phosphorylation of mTORC1 substrates in the skeletal muscle, signifying females may have higher mTORC1 activity (**Figs. 5H-J**). Interestingly, we found a striking decrease in phosphorylated S240/244 S6 and phosphorylated T37/S46 4E-BP1 in LP-fed Ovx mice relative to CTL-fed Ovx mice. In the case of phosphorylated S6, this recapitulates the effect of an LP diet on intact males. Notably, this ∼30-60% reduction in mTORC1 signaling was not observed in the liver of these same mice, demonstrating the potential tissue-specificity of this effect.

### LP-fed animals exhibit significant induction of iWAT beiging regardless of sex or gonadectomy

To visualize trends of the three experimental conditions on different phenotypic measures, we generated three different phenotypic heat maps isolating each experimental variable (**Fig. 6A**). While there are strong trends in diet effect across all groups, the males exhibited robust phenotypic responses to a LP diet that the female LP group does not. However, Ovx LP mice show a phenotypic pattern like that of the male LP group, as highlighted in the red-outlined boxes. Overall, gonadectomy significantly impacts body composition in both males and females with a reduction in lean mass and an increase in fat mass. As expected, we observed C57BL/6J female mice to be metabolically healthier than males in terms of phenotypic measures as shown in **Fig. 6A**.

**Figure 6.**
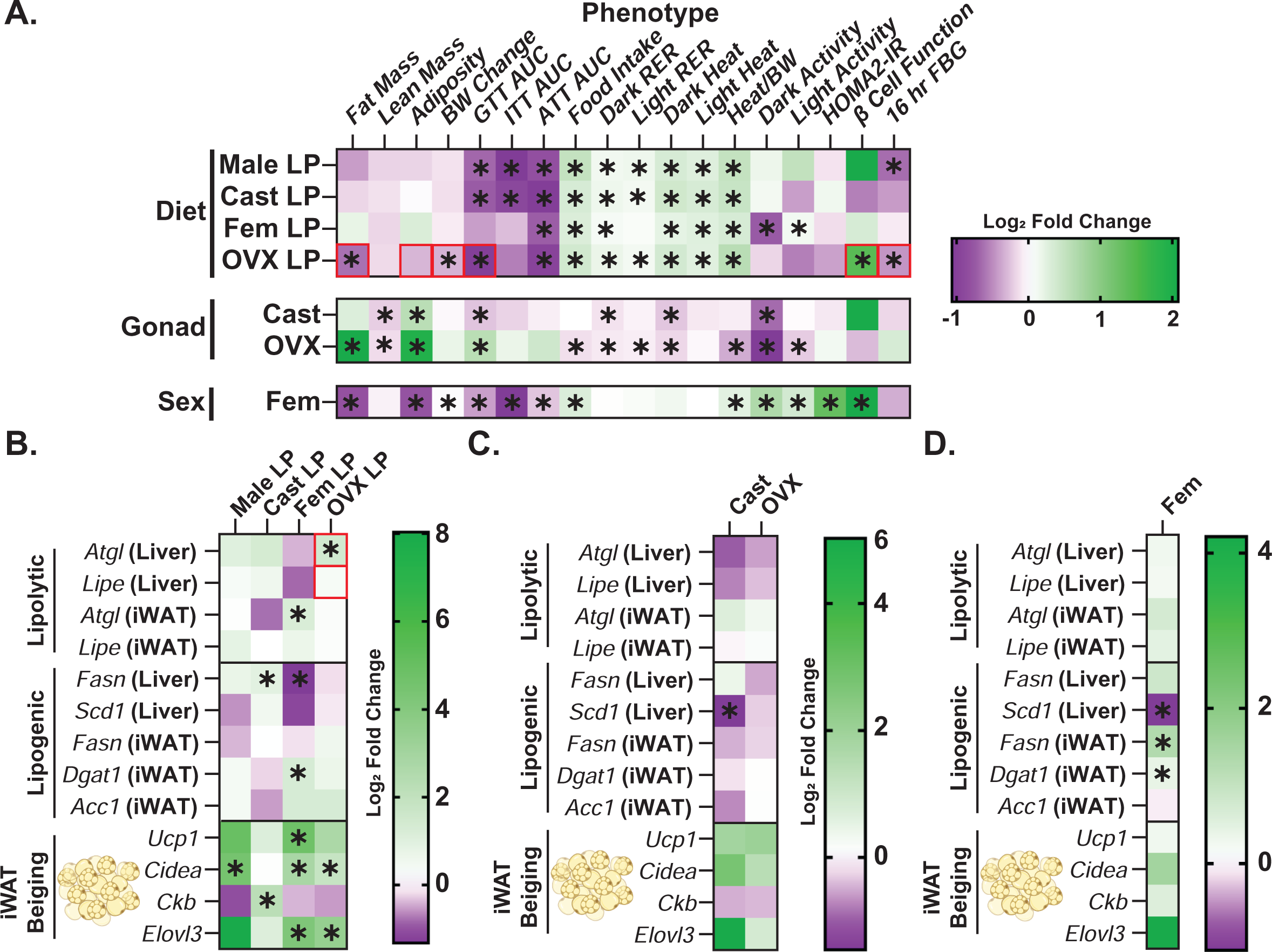
Altered lipid synthesis and breakdown promotes iWAT beiging that is sex, diet, and gonadectomy dependent. (A) Log2 fold changes of phenotypic measures isolating each experimental condition. (B-D) The Log2 fold change of liver and iWAT genes involved in lipid synthesis, fatty acid oxidation, and thermogenesis based on diet (B), gonadectomy (C), and sex (D). (A) n=11-12 mice per group. (B-D) n=4-6 mice per group. (B-D) The fold change was determined by comparing each group to its CTL diet (B), intact mice (C) or males (D). (A) Two-way ANOVA between sex, diet, or gonadectomy with post hoc Šidák’s adjusted test for pairwise comparisons, *p<0.05. p values for the overall effect of sex, diet, and gonadectomy and the interactions represent the significant p values from the two-way ANOVA. Data are represented as mean ±SEM. Abbreviations: CTL (21% control protein), LP (7% low protein), Cast (castration), Fem (female), Ovx (ovariectomy), Gonad (gonadectomy), BW (body weight), GTT (glucose tolerance test), AUC (area under the curve), ITT (insulin tolerance test), RER (respiratory exchange ratio), FBG (fasting blood glucose), iWAT (inguinal white adipose tissue).Created in BioRender. Knopf, B. (2026) https://BioRender.com/2yu9da3

As all LP-fed animals exhibited increased EE and FGF21 expression, we aimed to determine what other molecular processes could be driving the sex-dependent response to LP diets. To determine how lipid utilization was altered upon sex, diet, and gonadectomy, we analyzed genes involved in lipolysis, lipogenesis, and thermogenesis in the liver and iWAT, by qPCR (**Figs. S4A-M**). To summarize the results, we created heat maps for each experimental condition (**Figs. 6B-D**). Genes associated with lipolysis is reduced in the liver of female LP fed mice, whereas LP-fed OVX mice exhibit a similar mRNA expression profile of LP-fed males (**Fig. 6B**). Analysis of thermogenic genes revealed that consuming a LP diet induces beiging in the iWAT regardless of sex or surgical condition (**Fig. 6B**). Cast and Ovx alter lipid utilization in a tissue specific manner (**Fig. 6C**), as liver and iWAT exhibit different transcription profiles. We observed reduced hepatic lipolysis upon gonadectomy, but the effect of surgery on lipogenesis is unclear (**Fig. 6C**). We also found that gonadectomy itself activates iWAT beiging suggesting an important role for the sex organs in thermogenic regulation (**Fig. 6C**).

In **Fig. 6D**, we compared intact females to intact males on control diets to look at the differences in lipid metabolism between the two sexes. We see that females generally have increased lipolysis and lipogenesis in the liver and iWAT, however few are statistically significant. Lastly, we saw that females have increased expression of thermogenic genes compared to males, which may contribute to the altered metabolic response to a LP diet.

## Discussion

Dietary protein has clearly emerged as a critical regulator of metabolic health in rodents as well as in humans (Solon-Biet, McMahon et al. 2014, Solon-Biet, Mitchell et al. 2015). However, it is becoming clear that sex is an important factor in the response to dietary protein, with multiple studies showing that female mice are less responsive than male mice to a LP diet (Larson, Russo et al. 2017, Green, Pak et al. 2022). Here, we investigated the role of sex hormones in the sexually dimorphic response to LP diets in mice using a gonadectomy-based approach.

While intact females are resistant to LP diets, as have been previously shown, our results demonstrate that; Ovx makes female mice substantially more metabolically responsive. As previously reported (Larson, Russo et al. 2017), we observed a sexually dimorphic response of body weight as males consuming a LP diet have reduced body weight and females exhibit no change. Gonadectomy of both sexes resulted in increased fat accumulation that could not be reversed with LP in the males, but the fat accretion was significantly reduced by an LP diet in Ovx females. Similarly, a LP diet reduces body weight in Ovx mice that is not observed with intact females on LP, suggesting an interaction between ovarian products, fat accumulation, and dietary protein content. Analysis of glucose homeostasis revealed that the testes are not required for the LP induced improvements in glucose tolerance and insulin sensitivity. The Ovx mice on LP diet tend to mirror the male response through improved glucose tolerance and reduced fasting blood glucose levels.

At the molecular level, we characterized significant differences between intact animals and gonadectomized animals. We found that female mice have higher hepatic and circulating lipids-specifically TG. Interestingly we report that an LP diet further increases TG levels, although we observe no induction of *Gpt1*, a marker for liver damage. The increased TG storage in females may be explained by an overall increase of lipolysis and lipogenesis in female mice. We are just beginning to understand how the interaction between sex, gonadectomy, and dietary protein content alter lipid storage and utilization. Interestingly a LP diet and gonadectomy increases thermogenic gene expression in the iWAT of mice, however the induction of beiging in gonadectomy conditions does not equate to increased energy expenditure. FGF21 has long been thought to be a master puppeteer of increased energy expenditure and thus improved metabolic health. We found that FGF21 is robustly induced by an LP diet across all sexes and surgical conditions, suggesting that differences in the induction of FGF21 is not responsible for the blunted metabolic response of female mice. This data taken together highlights the importance of understanding how hormone signaling impacts energy homeostasis in the adipose tissue.

The data presented supports our hypothesis that removal of the ovaries permits female mice to metabolically respond to a LP diet. Female C57BL/6J mice reach puberty at 5 weeks of age and undergo cyclical fluctuations in sex hormones including estradiol. By 12 months of age, female mice reach reproductive senescence and estradiol levels remain low through the rest of their life. (Devillers, Mhaouty-Kodja et al. 2022). Since removal of the ovaries allows female mice to respond metabolically to PR, we suspect that female mice that are in reproductive senescence should respond as well. Indeed, late life protein restriction differs than early life restriction in that 22-month females on LP diets have improved glucose tolerance, although no changes in body composition were seen (Green, Pak et al. 2022, Yeh, Chini et al. 2024). Administering 17-β estradiol (E2) to ovariectomized mice rescues the phenotypes of increased accumulation of visceral adipose tissue, decreased glucose tolerance and worsened insulin sensitivity (Cruz, Santos et al. 2025). Our investigations in combination with studies utilizing exogenous E2 administration highlights the important interaction between E2 and metabolic pathways.

One of the limitations of this study is the timing at which gonadectomy was performed. We chose to gonadectomize mice at 5 weeks, which is prior to puberty based on previous research (Larson, Russo et al. 2017, Iwasa, Matsuzaki et al. 2018). The role of the sex organs, and specifically what hormones they produce, changes throughout lifespan. We speculate that gonadectomizing mice later in life will differentially impact metabolic processes compared to early life. Understanding how the timing of gonadectomy impacts metabolic health and aging in response to a low protein diet is an important future exploration, as previous studies show that gonadectomy timing can significantly impact late-life mortality in mice (Garratt, Try et al. 2021, Jiang, Cheng et al. 2023). Another limitation of this study is the utilization of only C57BL/6J mice. Performing these experiments in other mouse strains, especially those that are not genetically homogenous, could provide powerful insights into how gene variation contributes to these findings. Similarly, we examined the metabolic effect of removal of the sexual organs and the interaction with a low protein diet, but we did not examine which sex-specific hormone is responsible for the blunting effect of LP diets. Lastly, we performed tissue analysis on a limited number of tissues and molecular readouts. An unbiased multi-tissue omics approach could be beneficial in the future to determine differences between diet, sex, and gonadectomy.

Our findings not only highlight the important role for sex in the response to diet but also illuminate the gonads as a critical mediator of the response to dietary protein. We find, as have others in the past, that female mice are less metabolically responsive to a LP diet than male mice. Here, we identify the ovaries as critical mediators of this effect, showing that ovariectomy sensitizes female mice to the effects of a LP diet. If these results apply to humans, it could suggest that while young woman may not normally benefit from a LP diet, women who are postmenopausal or who have had their ovaries removed leading to metabolic deficits, might benefit metabolically from an LP diet. Further analysis into the role of sex hormones in the metabolic response to an LP diet will not only better complete the picture on how sex and sex organs impact the response to protein restriction but may also provide new insight on how the optimal human diet is affected by both sex and age.

## Data Availability Statement

The data that support the plots within this article and other findings of this study, including full scans of western blot images, are provided as Source Data files.

## Acknowledgements

We would like to thank all members of the Lamming lab for their assistance and input. The Lamming laboratory is supported in part by the NIH/NIA (AG056771, AG081482, AG084156, AG085898 and AG094153 to D.W.L.), NIH/NIDDK (DK125859 to D.W.L.) and startup funds from the University of Wisconsin-Madison School of Medicine and Public Health and Department of Medicine to D.W.L. M.F.C. was supported by F31AG082504. R.B. was supported by F31AG081115. C-Y.Y. was supported in part by a NIA F32 postdoctoral fellowship (F32AG077916), a NIA K99/R00 award (K99AG084921), and a Research Enrichment Component Postdoctoral Scholarship from the Wisconsin Alzheimer’s Disease Research Center (P30-AG062715). The authors utilized facilities and resources of the UW Carbone Cancer Center Experimental Animal Pathology Laboratory (P30CA014520). D.W.L. is a member of the Wisconsin Nathan Shock Center of Excellence in the Basic Biology of Aging, P30 AG092586. The Lamming lab was supported in part by the U.S. Department of Veterans Affairs (IS1-BX005524), and this work was supported using facilities and resources from the William S. Middleton Memorial Veterans Hospital. The content is solely the responsibility of the authors and does not necessarily represent the official views of the NIH. This work does not represent the views of the Department of Veterans Affairs or the United States Government.

## Author Contributions

B.A.K and D.W.L conceived and designed the experiments. B.A.K, I.G, B.A, T.R, M.M.S, M.C, R.B, Y.L, F.X, and C-Y.Y performed the experiments. B.A.K, B.A, T.R, M.M.S, M.C, R.B and D.W.L analyzed the data. B.A.K and D.W.L wrote the manuscript.

## References

Arriola Apelo, S. I., A. Lin, J. A. Brinkman, E. Meyer, M. Morrison, J. L. Tomasiewicz, C. P. Pumper, E. L. Baar, N. E. Richardson, M. Alotaibi and D. W. Lamming (2020). “Ovariectomy uncouples lifespan from metabolic health and reveals a sex-hormone-dependent role of hepatic mTORC2 in aging.” Elife 9.

Arvidsson Kvissberg, M. E., G. Hu, L. Chi, C. Bourdon, C. Ling, Y. ChenMi, K. Germain, I. P. van Peppel, L. Weise, L. Zhang, V. Di Giovanni, N. Swain, J. W. Jonker, P. Kim and R. Bandsma (2022). “Inhibition of mTOR improves malnutrition induced hepatic metabolic dysfunction.” Sci Rep 12(1): 19948.

Baar, E. L., K. A. Carbajal, I. M. Ong and D. W. Lamming (2016). “Sex- and tissue-specific changes in mTOR signaling with age in C57BL/6J mice.” Aging Cell 15(1): 155–166.

Calubag, M. F., I. Ademi, C. L. Green, H. S. M. Jayarathne, D. N. H. Manchanayake, S. M. Le, P. Lialios, L. E. Breuer, S. Yakar, R. Babygirija, M. M. Sonsalla, I. Grunow, C. Y. Yeh, Y. Liu, B. A. Knopf, W. A. Ricke, T. T. Liu, M. Sadagurski and D. W. Lamming (2025). “Lifelong restriction of dietary valine has sex-specific benefits for health and lifespan in mice.” bioRxiv.

Cruz, A. G. D., J. Santos, E. D. S. Alves, A. Santos, B. F. Trinca, F. N. Camargo, G. F. Bovolin and J. P. Camporez (2025). “Metabolic effects of late-onset estradiol replacement in high-fat-fed ovariectomized mice.” Curr Res Physiol 8: 100144.

Devillers, M. M., S. Mhaouty-Kodja and C. J. Guigon (2022). “Deciphering the Roles & Regulation of Estradiol Signaling during Female Mini-Puberty: Insights from Mouse Models.” Int J Mol Sci 23(22).

Ferraz-Bannitz, R., R. A. Beraldo, A. A. Peluso, M. Dall, P. Babaei, R. C. Foglietti, L. M. Martins, P. M. Gomes, J. S. Marchini, V. M. M. Suen, L. C. C. de Freitas, L. C. Navegantes, M. A. M. Pretti, M. Boroni, J. T. Treebak, M. A. Mori, M. C. Foss and M. C. Foss-Freitas (2022). “Dietary Protein Restriction Improves Metabolic Dysfunction in Patients with Metabolic Syndrome in a Randomized, Controlled Trial.” Nutrients 14(13).

Fontana, L., N. E. Cummings, S. I. Arriola Apelo, J. C. Neuman, I. Kasza, B. A. Schmidt, E. Cava, F. Spelta, V. Tosti, F. A. Syed, E. L. Baar, N. Veronese, S. E. Cottrell, R. J. Fenske, B. Bertozzi, H. K. Brar, T. Pietka, A. D. Bullock, R. S. Figenshau, G. L. Andriole, M. J. Merrins, C. M. Alexander, M. E. Kimple and D. W. Lamming (2016). “Decreased Consumption of Branched-Chain Amino Acids Improves Metabolic Health.” Cell Rep 16(2): 520–530.

Garratt, M., B. Bower, G. G. Garcia and R. A. Miller (2017). “Sex differences in lifespan extension with acarbose and 17-alpha estradiol: gonadal hormones underlie male-specific improvements in glucose tolerance and mTORC2 signaling.” Aging Cell 16(6): 1256–1266.

Garratt, M., K. A. Lagerborg, Y. M. Tsai, A. Galecki, M. Jain and R. A. Miller (2018). “Male lifespan extension with 17-alpha estradiol is linked to a sex-specific metabolomic response modulated by gonadal hormones in mice.” Aging Cell 17(4): e12786.

Garratt, M., H. Try and R. C. Brooks (2021). “Access to females and early life castration individually extend maximal but not median lifespan in male mice.” Geroscience 43(3): 1437–1446.

Green, C. L., D. W. Lamming and L. Fontana (2022). “Molecular mechanisms of dietary restriction promoting health and longevity.” Nat Rev Mol Cell Biol 23(1): 56–73.

Green, C. L., H. H. Pak, N. E. Richardson, V. Flores, D. Yu, J. L. Tomasiewicz, S. N. Dumas, K. Kredell, J. W. Fan, C. Kirsh, K. Chaiyakul, M. E. Murphy, R. Babygirija, G. A. Barrett-Wilt, J. Rabinowitz, I. M. Ong, C. Jang, J. Simcox and D. W. Lamming (2022). “Sex and genetic background define the metabolic, physiologic, and molecular response to protein restriction.” Cell Metab 34(2): 209–226 e205.

Harputlugil, E., C. Hine, D. Vargas, L. Robertson, B. D. Manning and J. R. Mitchell (2014). “The TSC complex is required for the benefits of dietary protein restriction on stress resistance in vivo.” Cell Rep 8(4): 1160–1170.

Hill, C. M., D. C. Albarado, L. G. Coco, R. A. Spann, M. S. Khan, E. Qualls-Creekmore, D. H. Burk, S. J. Burke, J. J. Collier, S. Yu, D. H. McDougal, H. R. Berthoud, H. Munzberg, A. Bartke and C. D. Morrison (2022). “FGF21 is required for protein restriction to extend lifespan and improve metabolic health in male mice.” Nat Commun 13(1): 1897.

Hill, C. M., T. Laeger, D. C. Albarado, D. H. McDougal, H. R. Berthoud, H. Munzberg and C. D. Morrison (2017). “Low protein-induced increases in FGF21 drive UCP1-dependent metabolic but not thermoregulatory endpoints.” Sci Rep 7(1): 8209.

Iwasa, T., T. Matsuzaki, K. Yano and M. Irahara (2018). “The effects of ovariectomy and lifelong high-fat diet consumption on body weight, appetite, and lifespan in female rats.” Horm Behav 97: 25–30.

Jiang, N., C. J. Cheng, J. Gelfond, R. Strong, V. Diaz and J. F. Nelson (2023). “Prepubertal castration eliminates sex differences in lifespan and growth trajectories in genetically heterogeneous mice.” Aging Cell 22(8): e13891.

Josse, J., Husson, François. (2016). missMDA: A Package for Handling Missing Values in Multivariate Data Analysis. Journal of Stastical Software.

Kalafut, K. C., Y. Cormerais, M. Y. Cisse, S. C. Lapp, K. E. Inouye, G. S. Hotamisligil and B. D. Manning (2025). “Dietary protein governs the role of insulin signaling in the postprandial regulation of hepatic mTORC1.” bioRxiv.

Kassambara, A. a. M., F. (2020). “Factoextra: Extract and Visualize the Results of Multivariate Data Analyses.”

Laeger, T., T. M. Henagan, D. C. Albarado, L. M. Redman, G. A. Bray, R. C. Noland, H. Munzberg, S. M. Hutson, T. W. Gettys, M. W. Schwartz and C. D. Morrison (2014). “FGF21 is an endocrine signal of protein restriction.” J Clin Invest 124(9): 3913–3922.

Lamming, D. W., N. E. Cummings, A. L. Rastelli, F. Gao, E. Cava, B. Bertozzi, F. Spelta, R. Pili and L. Fontana (2015). “Restriction of dietary protein decreases mTORC1 in tumors and somatic tissues of a tumor-bearing mouse xenograft model.” Oncotarget 6(31): 31233–31240.

Larson, K. R., K. A. Russo, Y. Fang, N. Mohajerani, M. L. Goodson and K. K. Ryan (2017). “Sex Differences in the Hormonal and Metabolic Response to Dietary Protein Dilution.” Endocrinology 158(10): 3477–3487.

Lê, S., Josse, Julie., Husson, François. (2008). “FactoMineR: An R Package for Multivariate Analysis.” Journal of Statistical Software 25(1).

Levine, M. E., J. A. Suarez, S. Brandhorst, P. Balasubramanian, C. W. Cheng, F. Madia, L. Fontana, M. G. Mirisola, J. Guevara-Aguirre, J. Wan, G. Passarino, B. K. Kennedy, M. Wei, P. Cohen, E. M. Crimmins and V. D. Longo (2014). “Low protein intake is associated with a major reduction in IGF-1, cancer, and overall mortality in the 65 and younger but not older population.” Cell Metab 19(3): 407–417.

Levy, J. C., D. R. Matthews and M. P. Hermans (1998). “Correct homeostasis model assessment (HOMA) evaluation uses the computer program.” Diabetes Care 21(12): 2191–2192.

Maida, A., A. Zota, K. A. Sjoberg, J. Schumacher, T. P. Sijmonsma, A. Pfenninger, M. M. Christensen, T. Gantert, J. Fuhrmeister, U. Rothermel, D. Schmoll, M. Heikenwalder, J. L. Iovanna, K. Stemmer, B. Kiens, S. Herzig and A. J. Rose (2016). “A liver stress-endocrine nexus promotes metabolic integrity during dietary protein dilution.” J Clin Invest 126(9): 3263–3278.

Mair, W., M. D. Piper and L. Partridge (2005). “Calories do not explain extension of life span by dietary restriction in Drosophila.” PLoS Biol 3(7): e223.

Mutel, E., A. Gautier-Stein, A. Abdul-Wahed, M. Amigo-Correig, C. Zitoun, A. Stefanutti, I. Houberdon, J. A. Tourette, G. Mithieux and F. Rajas (2011). “Control of blood glucose in the absence of hepatic glucose production during prolonged fasting in mice: induction of renal and intestinal gluconeogenesis by glucagon.” Diabetes 60(12): 3121–3131.

Nations, W. H. O. U. (2007). Protein and Amino Acid Requirements in Human Nutrition. Geneva: World Health Organization.

Ni Lochlainn, M., R. C. E. Bowyer, A. A. Welch, K. Whelan and C. J. Steves (2023). “Higher dietary protein intake is associated with sarcopenia in older British twins.” Age Ageing 52(2).

Nicolaisen, T. S., A. E. Lyster, K. A. Sjoberg, D. T. Haas, C. T. Voldstedlund, A. M. Lundsgaard, J. K. Jensen, E. M. Madsen, C. K. Nielsen, M. Bloch-Ibenfeldt, N. J. Wewer Albrechtsen, A. J. Rose, N. Krahmer, C. Clemmensen, E. A. Richter, A. M. Fritzen and B. Kiens (2025). “Dietary protein restriction elevates FGF21 levels and energy requirements to maintain body weight in lean men.” Nat Metab 7(3): 602–616.

Okun, J. G., P. M. Rusu, A. Y. Chan, Y. Wu, Y. W. Yap, T. Sharkie, J. Schumacher, K. V. Schmidt, K. M. Roberts-Thomson, R. D. Russell, A. Zota, S. Hille, A. Jungmann, L. Maggi, Y. Lee, M. Bluher, S. Herzig, M. A. Keske, M. Heikenwalder, O. J. Muller and A. J. Rose (2021). “Liver alanine catabolism promotes skeletal muscle atrophy and hyperglycaemia in type 2 diabetes.” Nat Metab 3(3): 394–409.

Richardson, N. E., E. N. Konon, H. S. Schuster, A. T. Mitchell, C. Boyle, A. C. Rodgers, M. Finke, L. R. Haider, D. Yu, V. Flores, H. H. Pak, S. Ahmad, S. Ahmed, A. Radcliff, J. Wu, E. M. Williams, L. Abdi, D. S. Sherman, T. Hacker and D. W. Lamming (2021). “Lifelong restriction of dietary branched-chain amino acids has sex-specific benefits for frailty and lifespan in mice.” Nat Aging 1(1): 73–86.

Simonson, D. C. and R. A. DeFronzo (1990). “Indirect calorimetry: methodological and interpretative problems.” Am J Physiol 258(3 Pt 1): E399–412.

Sluijs, I., J. W. Beulens, A. D. van der, A. M. Spijkerman, D. E. Grobbee and Y. T. van der Schouw (2010). “Dietary intake of total, animal, and vegetable protein and risk of type 2 diabetes in the European Prospective Investigation into Cancer and Nutrition (EPIC)-NL study.” Diabetes Care 33(1): 43–48.

Solon-Biet, S. M., A. C. McMahon, J. W. Ballard, K. Ruohonen, L. E. Wu, V. C. Cogger, A. Warren, X. Huang, N. Pichaud, R. G. Melvin, R. Gokarn, M. Khalil, N. Turner, G. J. Cooney, D. A. Sinclair, D. Raubenheimer, D. G. Le Couteur and S. J. Simpson (2014). “The ratio of macronutrients, not caloric intake, dictates cardiometabolic health, aging, and longevity in ad libitum-fed mice.” Cell Metab 19(3): 418–430.

Solon-Biet, S. M., S. J. Mitchell, S. C. Coogan, V. C. Cogger, R. Gokarn, A. C. McMahon, D. Raubenheimer, R. de Cabo, S. J. Simpson and D. G. Le Couteur (2015). “Dietary Protein to Carbohydrate Ratio and Caloric Restriction: Comparing Metabolic Outcomes in Mice.” Cell Rep 11(10): 1529–1534.

Sookoian, S., G. O. Castano, R. Scian, T. Fernandez Gianotti, H. Dopazo, C. Rohr, G. Gaj, J. San Martino, I. Sevic, D. Flichman and C. J. Pirola (2016). “Serum aminotransferases in nonalcoholic fatty liver disease are a signature of liver metabolic perturbations at the amino acid and Krebs cycle level.” Am J Clin Nutr 103(2): 422–434.

Wei, T., Simko, Viliam., Levy, Michael., Xie, Yihui., Jin, Yan., Zemla, Jeff., Freidank, Moritz., Cai, Jun., Protivinsky, Tomas. (2024). “corrplot: Visualization of a Correlation Matrix.”

Yeh, C. Y., L. C. S. Chini, J. W. Davidson, G. G. Garcia, M. S. Gallagher, I. T. Freichels, M. F. Calubag, A. C. Rodgers, C. L. Green, R. Babygirija, M. M. Sonsalla, H. H. Pak, M. E. Trautman, T. A. Hacker, R. A. Miller, J. A. Simcox and D. W. Lamming (2024). “Late-life protein or isoleucine restriction impacts physiological and molecular signatures of aging.” Nat Aging 4(12): 1760–1771.

