## Supplemental Figures for "Female resistance to the metabolic benefits of protein restriction is reversed by ovariectomy in mice"

### Figure S1

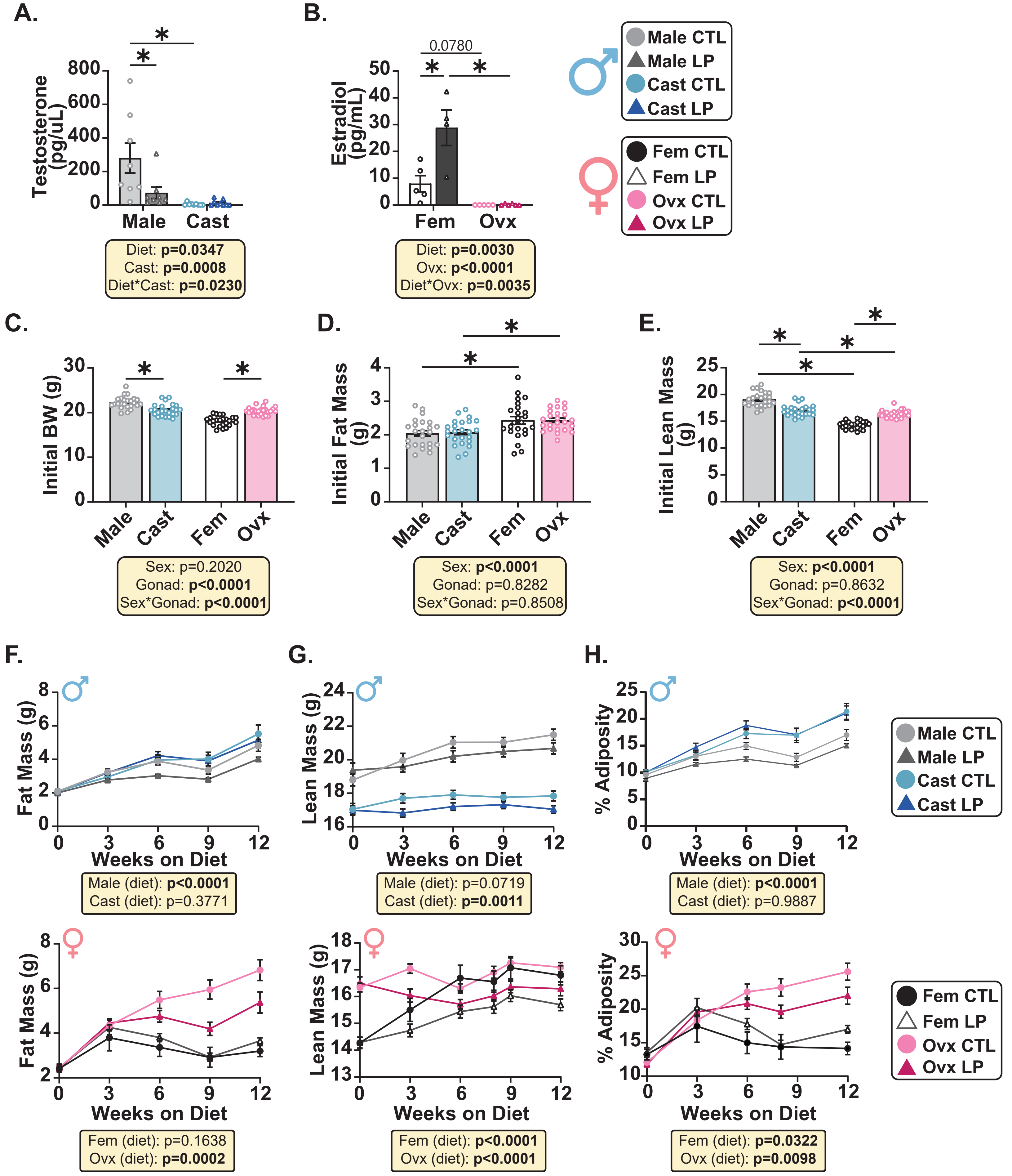
A.
Testosterone (pg/uL)
Male
Cast
Diet: p=0.0347
Cast: p=0.0008
Diet*Cast: p=0.0230
B.
Estradiol (pg/mL)
0.0780
Fem
Ovx
Diet: p=0.0030
Ovx: p<0.0001
Diet*Ovx: p=0.0035
Male CTL
Male LP
Cast CTL
Cast LP
Fem CTL
Fem LP
Ovx CTL
Ovx LP
C.
Initial BW (g)
Male
Cast
Fem
Ovx
Sex: p=0.2020
Gonad: p<0.0001
Sex*Gonad: p<0.0001
D.
Initial Fat Mass (g)
Sex: p<0.0001
Gonad: p=0.8282
Sex*Gonad: p=0.8508
E.
Initial Lean Mass (g)
Sex: p<0.0001
Gonad: p=0.8632
Sex*Gonad: p<0.0001
F.
Fat Mass (g)
Weeks on Diet
Male (diet): p<0.0001
Cast (diet): p=0.3771
G.
Lean Mass (g)
Weeks on Diet
Male (diet): p=0.0719
Cast (diet): p=0.0011
H.
% Adiposity
Weeks on Diet
Male (diet): p<0.0001
Cast (diet): p=0.9887
Fat Mass (g)
Weeks on Diet
Fem (diet): p=0.1638
Ovx (diet): p=0.0002
Lean Mass (g)
Weeks on Diet
Fem (diet): p<0.0001
Ovx (diet): p<0.0001
% Adiposity
Weeks on Diet
Fem (diet): p=0.0322
Ovx (diet): p=0.0098

**Supplemental Figure 1. Gonadectomy alters body composition throughout the 12-week study.**

Quantification of circulating testosterone (A) and estradiol (B) using refed plasma by ELISA. (C-E) Characterization of body weight (C), lean mass (D), and fat mass (E) prior to beginning the experiment. (F-H) Time dependent changes in fat mass (F), lean mass (G), and adiposity (H) during the 12-week period. (A-B) 5-8 mice per group. (C-H) n=11-12 mice per group. (A-B, F-H) Two-way ANOVA between diet and gonadectomy. Statistics for the overall effects of diet, gonadectomy, and the interaction represent the p value from a two-way ANOVA. *$p<0.05$ from a Šidák's post-test examining the effect of parameters identified as significant in the two-way ANOVA. (C-E) Two-way ANOVA between sex and gonadectomy with post hoc Šidák's adjusted test for pairwise comparisons, *$p<0.05$. p values for the overall effect of sex and gonadectomy and the interactions represent the significant p values from the two-way ANOVA. Data are represented as mean ±SEM Abbreviations: CTL (21% control protein diet), LP (7% low protein diet), Cast (castration), Fem (female), Ovx (ovariectomy).

### Figure S2

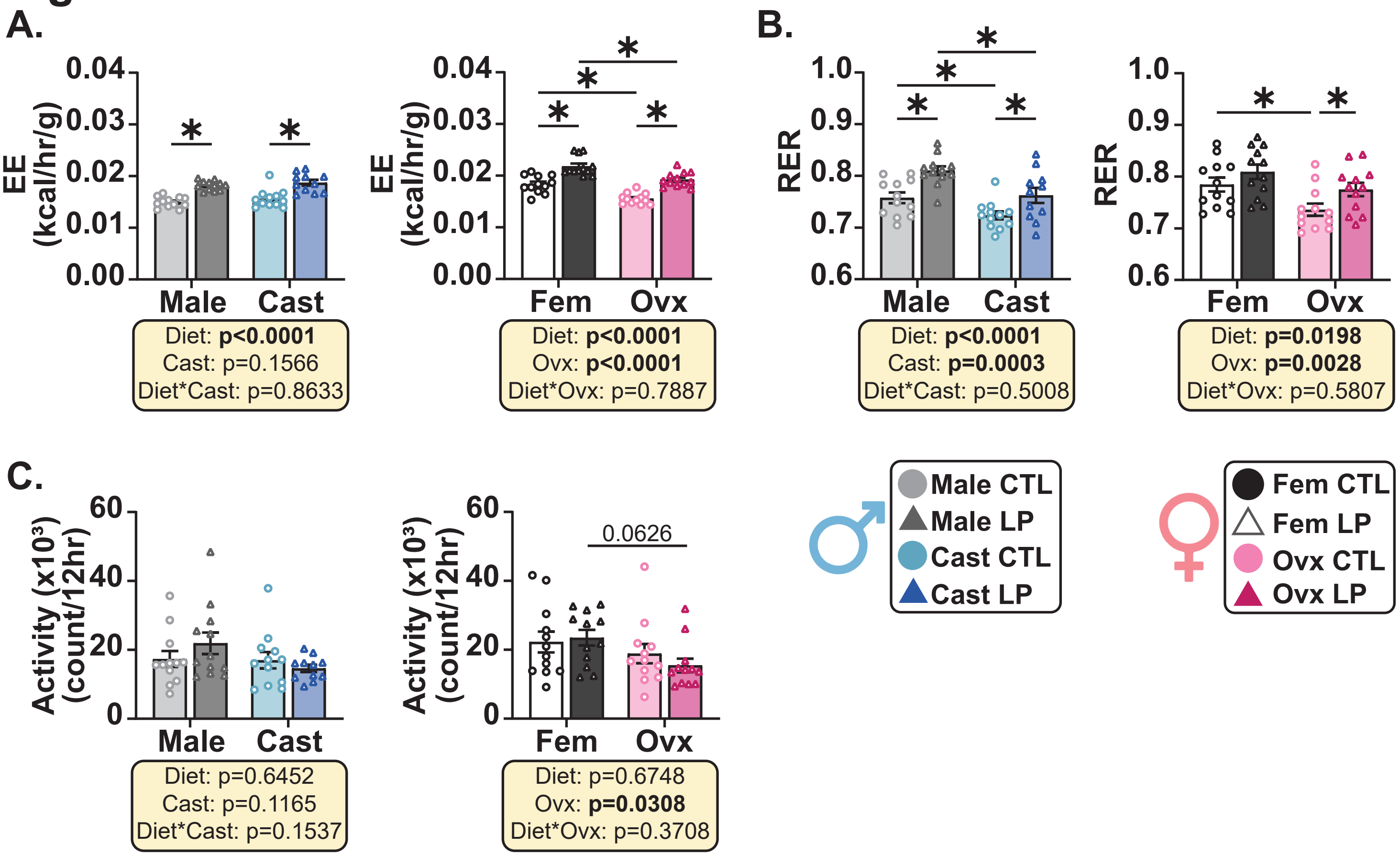

A.
EE (kcal/hr/g)
0.04
0.03
0.02
0.01
0.00
Male
Cast
Diet: **p<0.0001**
Cast: p=0.1566
Diet*Cast: p=0.8633
Fem
Ovx
Diet: **p<0.0001**
Ovx: **p<0.0001**
Diet*Ovx: p=0.7887
B.
RER
1.0
0.9
0.8
0.7
0.6
Diet: **p<0.0001**
Cast: **p=0.0003**
Diet*Cast: p=0.5008
Diet: **p=0.0198**
Ovx: **p=0.0028**
Diet*Ovx: p=0.5807
C.
Activity (x10³) (count/12hr)
60
40
20
0
Diet: p=0.6452
Cast: p=0.1165
Diet*Cast: p=0.1537
0.0626
Diet: p=0.6748
Ovx: **p=0.0308**
Diet*Ovx: p=0.3708
Male CTL
Male LP
Cast CTL
Cast LP
Fem CTL
Fem LP
Ovx CTL
Ovx LP

**Supplemental Figure 2. Dietary protein content dominates changes in energy balance during the light cycle.**

(A) Average Energy Expenditure (EE) normalized to body weight during the light cycle. (B) Average RER during the 12-hour light cycle. (C) Average spontaneous activity during the light cycle. (A-C) n=11-12 mice/group. (A-C) Statistics for the overall effects of diet, gonadectomy, and the interaction represent the p value from a two-way ANOVA. *$p<0.05$ from a Šidák's post-test examining the effect of parameters identified as significant in the two-way ANOVA. Data are represented as mean ±SEM. Abbreviations: CTL (21% control protein diet), LP (7% low protein diet), Cast (castration), Fem (female), Ovx (ovariectomy), EE (energy expenditure), RER (respiratory exchange ratio).

### Figure S3

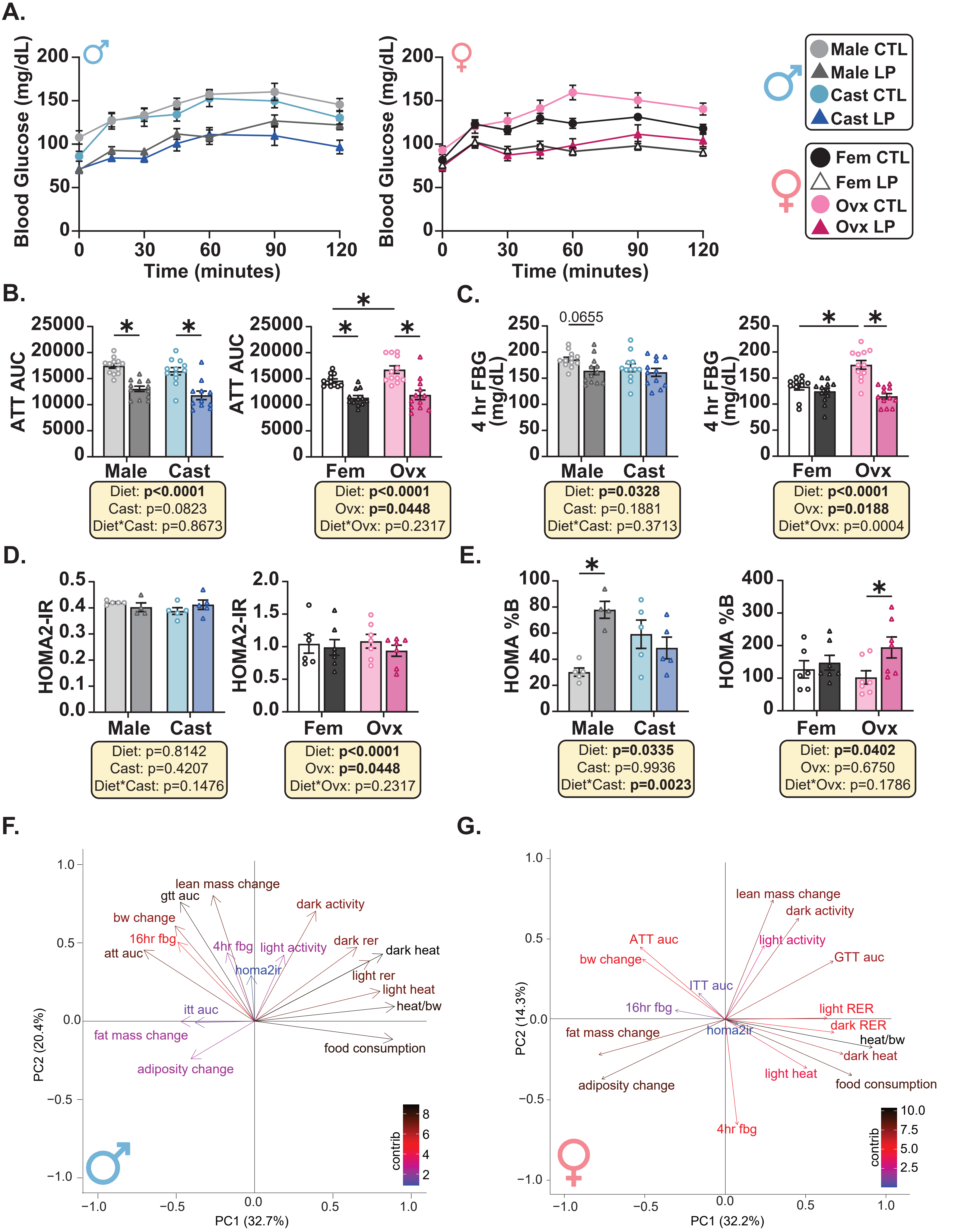

A.
Blood Glucose (mg/dL)
Time (minutes)
Male CTL
Male LP
Cast CTL
Cast LP
Fem CTL
Fem LP
Ovx CTL
Ovx LP
B.
ATT AUC
Male Cast
Diet: p<0.0001
Cast: p=0.0823
Diet*Cast: p=0.8673
Fem Ovx
Diet: p<0.0001
Ovx: p=0.0448
Diet*Ovx: p=0.2317
C.
4 hr FBG (mg/dL)
0.0655
Male Cast
Diet: p=0.0328
Cast: p=0.1881
Diet*Cast: p=0.3713
Fem Ovx
Diet: p<0.0001
Ovx: p=0.0188
Diet*Ovx: p=0.0004
D.
HOMA2-IR
Male Cast
Diet: p=0.8142
Cast: p=0.4207
Diet*Cast: p=0.1476
Fem Ovx
Diet: p<0.0001
Ovx: p=0.0448
Diet*Ovx: p=0.2317
E.
HOMA %B
Male Cast
Diet: p=0.0335
Cast: p=0.9936
Diet*Cast: p=0.0023
Fem Ovx
Diet: p=0.0402
Ovx: p=0.6750
Diet*Ovx: p=0.1786
F.
lean mass change
gtt auc
dark activity
bw change
16hr fbg
4hr fbg
light activity
dark rer
att auc
dark heat
homa2ir
light rer
light heat
itt auc
heat/bw
fat mass change
food consumption
adiposity change
contrib
PC1 (32.7%)
PC2 (20.4%)
G.
lean mass change
dark activity
ATT auc
light activity
bw change
GTT auc
ITT auc
16hr fbg
light RER
dark RER
fat mass change
homa2ir
heat/bw
dark heat
light heat
adiposity change
food consumption
4hr fbg
contrib
PC1 (32.2%)
PC2 (14.3%)

**Supplemental Figure 3. Sex, diet, and gonadectomy impact insulin signaling and metabolic pathways.**

(A-B) Alanine tolerance test (ATT) and area under the curve (AUC) calculation after 5 weeks of dietary intervention. (C) Fasting blood glucose (FBG) values after a 4-hour fast. (D-E) HOMA2-IR calculation and β-Cell function (E) based on fasting insulin and blood glucose levels. (F-G) Contribution plot of variables to PC1 and PC2 for males (F) and females (G). (A-C, F-G) n=11-12 mice/group. (D-E) n=3-7 mice per group. (B-E) Statistics for the overall effects of diet, gonadectomy, and the interaction represent the p value from a two-way ANOVA. *$p<0.05$ from a Šidák's post-test examining the effect of parameters identified as significant in the two-way ANOVA. Data are represented as mean ±SEM. Abbreviations: CTL (21% control protein diet), LP (7% low protein diet), Cast (castration), Fem (female), Ovx (ovariectomy), BW (body weight), GTT (glucose tolerance test), ITT (insulin tolerance test), RER (respiratory exchange ratio), FC (food consumption), ATT (alanine tolerance test), AUC (area under the curve), FBG (fasting blood glucose).

### Figure S4

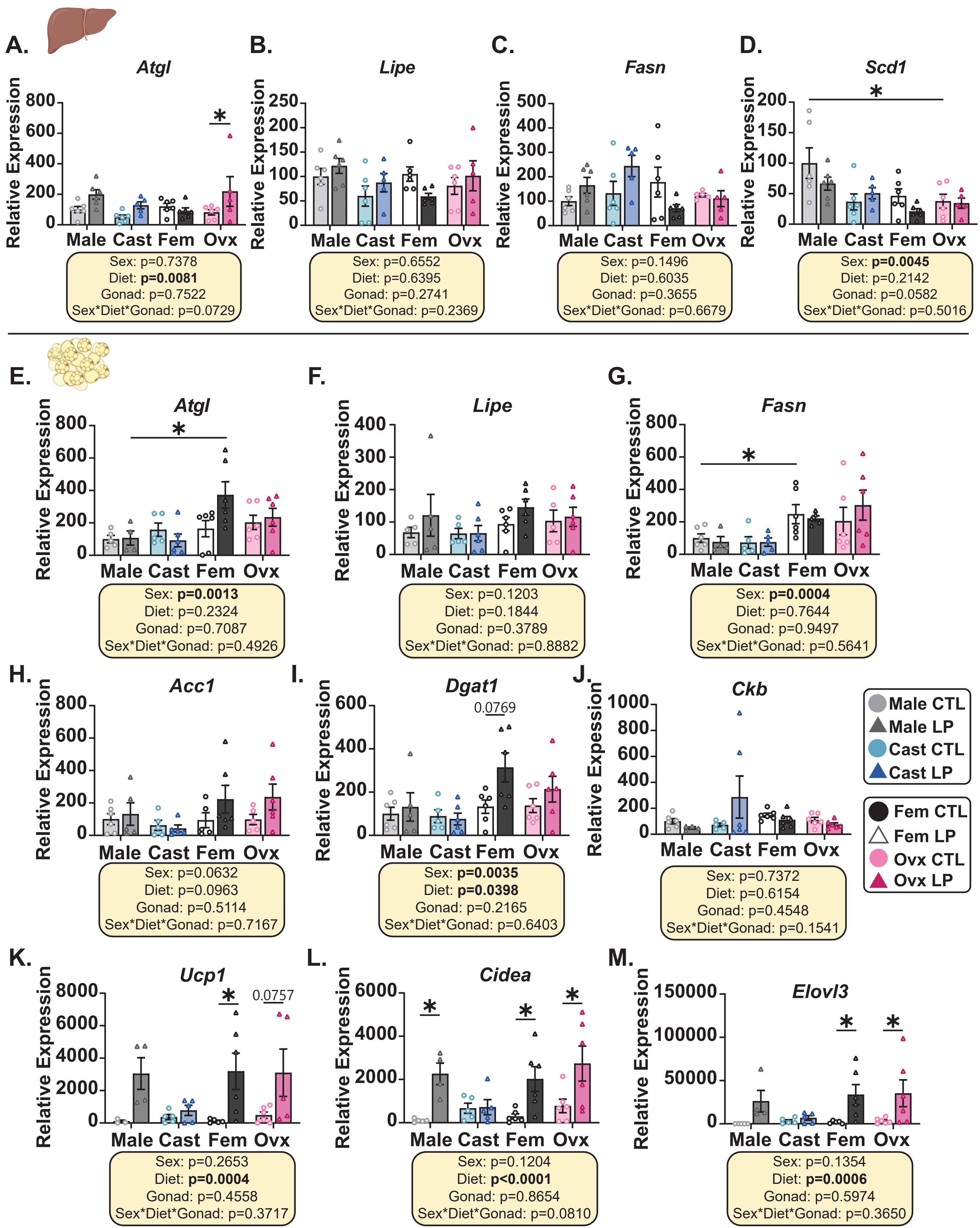
A.
Atgl
Relative Expression
Male Cast Fem Ovx
Sex: p=0.7378
Diet: p=0.0081
Gonad: p=0.7522
Sex*Diet*Gonad: p=0.0729
B.
Lipe
Sex: p=0.6552
Diet: p=0.6395
Gonad: p=0.2741
Sex*Diet*Gonad: p=0.2369
C.
Fasn
Sex: p=0.1496
Diet: p=0.6035
Gonad: p=0.3655
Sex*Diet*Gonad: p=0.6679
D.
Scd1
Sex: p=0.0045
Diet: p=0.2142
Gonad: p=0.0582
Sex*Diet*Gonad: p=0.5016
E.
Atgl
Sex: p=0.0013
Diet: p=0.2324
Gonad: p=0.7087
Sex*Diet*Gonad: p=0.4926
F.
Lipe
Sex: p=0.1203
Diet: p=0.1844
Gonad: p=0.3789
Sex*Diet*Gonad: p=0.8882
G.
Fasn
Sex: p=0.0004
Diet: p=0.7644
Gonad: p=0.9497
Sex*Diet*Gonad: p=0.5641
H.
Acc1
Sex: p=0.0632
Diet: p=0.0963
Gonad: p=0.5114
Sex*Diet*Gonad: p=0.7167
I.
Dgat1
0.0769
Sex: p=0.0035
Diet: p=0.0398
Gonad: p=0.2165
Sex*Diet*Gonad: p=0.6403
J.
Ckb
Relative Expession
Sex: p=0.7372
Diet: p=0.6154
Gonad: p=0.4548
Sex*Diet*Gonad: p=0.1541
Male CTL
Male LP
Cast CTL
Cast LP
Fem CTL
Fem LP
Ovx CTL
Ovx LP
K.
Ucp1
0.0757
Sex: p=0.2653
Diet: p=0.0004
Gonad: p=0.4558
Sex*Diet*Gonad: p=0.3717
L.
Cidea
Sex: p=0.1204
Diet: p<0.0001
Gonad: p=0.8654
Sex*Diet*Gonad: p=0.0810
M.
Elovl3
Sex: p=0.1354
Diet: p=0.0006
Gonad: p=0.5974
Sex*Diet*Gonad: p=0.3650

**Supplemental Figure 4. LP diets increase lipolysis, lipogenesis, and thermogenesis in a sex, gonadectomy, and tissue specific manner.**

(A-D) mRNA expression of *Atgl* (A), *Lipe* (B), *Fasn* (C), and *Scd1* (D) in liver relative to male CTL by qPCR analysis. (E-M) mRNA expression of *Atgl* (E), *Lipe* (F), *Fasn* (G), *Acc1* (H), *Dgat* (I), *Ckb* (J), *Ucp1* (K), *Cidea* (L), and *Elovl3* (M) in iWAT relative to male CTL by qPCR analysis. (A-M) n=4-6 mice per group. (A-C) Three-way ANOVA between sex, diet, and gonadectomy with post hoc Šidák's adjusted test for pairwise comparisons, *$p<0.05$. p values for the overall effect of sex, diet, and gonadectomy and the interactions represent the significant p values from the three-way ANOVA. Abbreviations: CTL (21% control protein diet), LP (7% low protein diet), Cast (castration), Fem (female), Ovx (ovariectomy), Gonad (gonadectomy). Created in BioRender. Knopf, B (2026). https://BioRender.com/n3hbix1
